# Phase-dependent closed-loop intersectional short-pulse stimulation reduces seizure duration: From computational modeling to clinical application

**DOI:** 10.64898/2026.08.19.745828

**Authors:** Lívia Barcsai, Nóra Forgó, Zoltán Somogyvári, Veronika Házi, Kristóf Furuglyás, Márton Huszár-Kis, Zoltán Chadaide, Péter Ráfi, Tamás Laszlovszky, Loránd Erőss, Antal Berényi

## Abstract

Drug-resistant epilepsy affects one-third of patients with persistent seizures despite optimal therapy. Intersectional short-pulse (ISP) stimulation is a novel transcranial electrical stimulation technique designed to deliver temporally precise, spatially targeted modulation of pathological brain activity. Here, we combined computational modeling with measurements in a rat epilepsy model and in patients with epilepsy to map the relationship between stimulation phase and seizure attenuation. In silico simulations of epileptiform networks showed that ISP stimulation significantly shortened seizure duration, with efficacy strongly depending on the phase of delivery. Phase-targeted stimulation during the rising phase and around the peaks (∼45–90°) of the seizure oscillations led to the greatest reduction in seizure length. In rodents, ISP decreased seizure duration by 42.4% and shortened generalized seizure segments by 58.3%. In humans, stimulation reduced seizure length by 60.9% compared to control seizures. Phase dependence was evident across models and species, with a prominent efficacy window in the rising-to-peak portion of the ictal oscillation and model-specific secondary windows. These findings show that phase-targeted ISP can substantially shorten seizures and support phase-resolved stimulation as a precision-neuromodulation approach for epilepsy.

## Introduction

Epilepsy is among the most prevalent neurological disorders worldwide, with an estimated 70 million people affected, underscoring its status as a major global health concern^1^. While many patients achieve long-term remission with treatment, up to one-third develop drug-resistant epilepsy, which is associated with poor quality of life, increased comorbidity, and higher mortality^2^. Epilepsy is not a single disease but a spectrum of disorders with diverse etiologies, network substrates and clinical outcomes^3,4^. This heterogeneity creates a major challenge for treatment: a universally effective intervention cannot be expected to act on a single upstream cause.

Despite this heterogeneity, seizures provide a common dynamical convergence point: neuronal populations enter abnormal, self-sustaining network states, commonly expressed as structured electrographic oscillations. Dynamical-systems analyses indicate that seizure onset, evolution and termination can be understood as transitions between network regimes, although individual seizures may belong to different dynamical classes^5,6^. Pathological ictal dynamics therefore do not represent one mechanism shared by all epilepsies; rather, they constitute a common, time-resolved intervention point downstream of diverse causes. Perturbing these evolving dynamics may therefore provide an intervention strategy that operates downstream of - and potentially across - distinct underlying etiologies^7–9^.

For patients who are not candidates for successful resective surgery, implantable neuromodulation offers an important but still incomplete alternative. Conventional vagus nerve stimulation (VNS) and anterior thalamic deep-brain stimulation (DBS) predominantly deliver stimulation according to programmed duty cycles, whereas responsive neurostimulation (RNS) detects abnormal intracranial activity and rapidly stimulates one or two predefined targets^10–13^. Although these approaches can produce substantial long-term reductions in seizure frequency, therapeutic responses remain heterogeneous and complete seizure control is uncommon. Moreover, event-triggered stimulation does not necessarily provide dynamical precision: current systems do not explicitly determine whether the evolving ictal network is momentarily susceptible to the delivered perturbation. This leaves a central control variable unresolved: when within the seizure dynamics should stimulation be applied to maximize its terminating effect?

Against this background, we previously demonstrated that closed-loop transcranial electrical stimulation (TES) can shorten epileptic seizures in animal models, showing that seizure-triggered interventions can selectively disrupt pathological brain activity while leaving other functions intact^14^. These experiments highlighted the potential of closed-loop neuromodulation but also raised questions about how stimulation could be made more effective and broadly translatable. Disrupting pathological seizure patterns with high fidelity requires intracerebral electrical gradients of approximately 1 mV/mm to directly influence neuronal spiking^14,15^. However, current shunting through the skull renders TES painful when it is applied at the intensities required for reliable seizure control^16^. Intersectional short-pulse (ISP) stimulation can achieve sufficiently strong intracerebral fields while keeping the scalp sensation tolerable by distributing ultra-brief, high-intensity electrical pulses across multiple electrode pairs^17,18^. ISP thus offers a technically feasible solution for non-invasive neuromodulation in epilepsy^16,18,19^.

Computational modeling combined with in vivo single-cell patch-clamp recordings showed that neurons integrate ISP- induced fields in a non-vectorial manner, enabling selective modulation of cortical and subcortical networks^18^. In rodent models of temporal lobe epilepsy, closed-loop ISP substantially shortened seizure duration and reduced severity, confirming its effectiveness in preclinical experiments^18^.

Closed-loop ISP stimulation through subgaleally implanted electrodes significantly reduced seizure duration and the likelihood of secondary generalization in a first-in-patient clinical study, while remaining safe and well tolerated^19^.

While closed-loop ISP stimulation has proven effective in shortening seizures, the role of stimulation timing within the seizure oscillation remains unresolved. Prior studies have suggested that the phase of the oscillatory cycle may critically influence the efficacy of stimulation^20–22^. However, systematic investigation of this principle has not been performed. The present study aimed to address this gap by introducing phase-targeted ISP stimulation as a refinement of closed-loop control. Computational modeling of epileptiform networks was first used to predict phase-dependent effects of ISP on seizure dynamics. These predictions were then validated in hippocampal-kindled rats with real-time phase-locked ISP stimulation, and finally tested in retrospective analysis of human recordings obtained with subgaleal electrodes. The overarching goal was to identify the oscillatory phase at which ISP stimulation most effectively shortens seizures, and to examine the extent to which phase-dependent response profiles are shared across models and species. By bridging modeling, preclinical experiments, and clinical data, this work advances ISP neuromodulation from general closed-loop seizure control toward precisely phase-targeted therapeutic intervention.

## Methods

### In silico modeling

Epileptic activity was simulated in a random network of 100 population models, each corresponding to a four-variable extended Morris–Lecar model^23^. Each population was described by four dynamical variables: V, W, Z and Ca2+ concentration, with calcium providing additional slow dynamics.

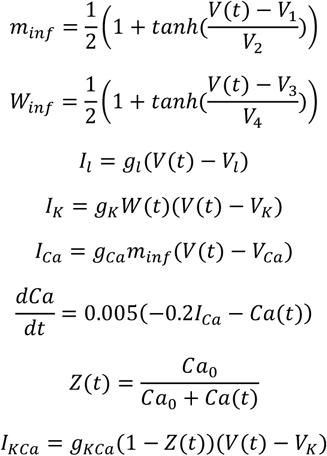

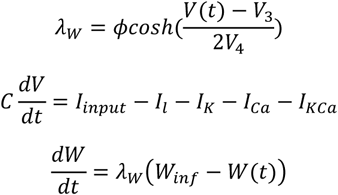

The parameter values were: VK = −84; Vl = −60; VCa = 120; gK = 8; gl = 2; gCa = 4; gKCa = 0.25; C = 20; V1 = −1.2; V2 = 18; V3 = 12; V4 = 17; φ = 0.04; Ca0 = 10; ɸ = 0.242;

The initial values were V_(t0)_ = −35, W_(t0)_ = 0.0148, Z_(t0)_ = 0.1 and Ca_(t0)_ = 10, and the simulations were integrated with a time step of dt = 0.1 ms.

The total input current to the j^th^ node consists of two components:

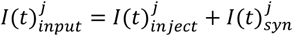

The synaptic current describing interactions within the network was: 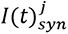

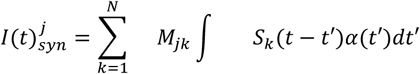

where M_jk_ denotes the jk^th^ element of the connection matrix M, and Sk(t) is a binary variable that equals 1 if node *k* emits a spike at time t and 0 otherwise. For each simulation, the binary connection matrix M was generated by independently setting each element to 1 with probability p = 0.1. Thus, the expected number of connections was 1000 in the 100 × 100 connection matrix.

Upon activation, each node generated a synaptic current described by an alpha function:

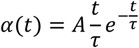

where A = 100 and *τ* = 10.

The background excitation was described as: 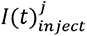

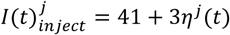

where η(t) is a random variable drawn from an independent normal distribution for all nodes. The system fluctuated most of the time but occasionally exhibited spontaneous seizure-like activity. However, during the simulation, at a specified time the injected current was changed to

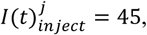

for a 100-ms period starting at t = 1 s. This initial stimulation elicited seizure-like activity in 87 out of 100 simulated cases.

The extracellular potential originates from net membrane current, comprising conductive transmembrane and capacitive components^24^. Because net membrane current sums to zero along an entire neuron, only spatial inhomogeneities can generate dipolar or higher-order extracellular fields. The Morris–Lecar model is a lumped, single-compartment point model and therefore cannot directly generate such spatial inhomogeneity. We therefore assumed that the modeled membrane currents were more abundant at one end of the neuron (i.e., the somatic end), allowing the extracellular potential to be approximated as a dipole field whose amplitude is proportional to the conductive membrane current and, consequently, to the derivative of membrane potential.

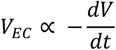

The sign of the proportionality depends on the spatial distribution of the currents on the neuron, the orientation of the neuron relative to the electrode, and the location of the reference electrode. Thus, the extracellular potential was approximated by the negative derivative of the population mean membrane potential.

To determine oscillatory phase, this approximation was low-pass filtered by convolution with a Gaussian sliding window (σ = 50 steps). The instantaneous phase was calculated as the angle of the analytic signal obtained by applying the Hilbert transform to the filtered extracellular potential*V_EC_*.

The 10-s simulations (100,000 steps) were repeated 100 times. For each simulation, a new connection matrix and a new set of random inputs η(t) were generated.

Epileptic activity was either started spontaneously or elicited by initial stimulation. After the initiation of the seizure, each simulation was run twice, in two distinct ways: first, it ran autonomously without further intervention; second, the phase-dependent simulated ISP stimulation, intended to stop the seizure, was initiated after 1.25 seconds of simulation time. ISP stimulation intensity was set to 500 based on an iterative dose-titration experiment (Fig. S1). The two simulations started from exactly the same system state and evolved deterministically; this procedure therefore enabled a matched comparison of seizure duration with and without stimulation, which is not possible in experimental recordings. The length of the seizure was defined as the time interval between the initiation and the last point where the mean membrane potential was above 0 mV.

### State-space reconstruction and dimension estimation

State-space reconstruction was performed using time-delay embedding^25^ applied to the population-mean membrane potential time series. The signal was temporally undersampled by a factor of 10 and embedded into a five-dimensional state space using a time delay of τ = 5 ms. In this embedding, epileptic activity formed a coherent oscillatory trajectory, while the resting state corresponded to a fixed point in state space.

Simulations were conducted on 100 randomly generated networks with random initial conditions and independent noise realizations. Of these simulations, 87 exhibited seizure durations exceeding 4 s and were included in the subsequent analysis. The intrinsic dimension of the reconstructed attractor was estimated using a k-nearest-neighbor–based method^26^, with the number of neighbors set to k = 10. Although the full system comprised 100 Morris–Lecar population models (400 dynamical variables), the reconstructed attractors exhibited a median dimension of 3.07±0.38 (median absolute deviation). The observed low dimensionality suggests that, during seizures, network activity and its normally high degrees of freedom become confined to an attractor resembling the structure of a Rössler attractor^27^, allowing seizure dynamics and stimulation effects to be studied in a reduced-dimensional representation.

### Animal experiments

#### Animals

Twenty-one young adult male rats were used (9 Wistar and 12 Long–Evans rats; 300–450 g). The animals were housed in groups of 2–3 per cage before surgery and individually afterward for the remainder of the experiment. The animals were maintained under controlled environmental conditions with a 12-hour light/dark cycle, constant temperature, and humidity. Rats had access to water and a standard commercial diet ad libitum throughout the study period. All experimental procedures were conducted according to the European Union guidelines (2003/65/CE) and the National Institutes of Health Guidelines for the Care and Use of Animals for Experimental Procedures. The study protocols received approval from the Ethical Committee for Animal Research at the Albert Szent-Györgyi Medical and Pharmaceutical Center of the University of Szeged (XIV/1248/2018 and XIV/824/2021).

#### Surgery

The surgical protocol has been previously described in detail by Kozák et al. in 2018 ^28^. Briefly, the rats were anesthetized with 2% isoflurane and head-fixed in the stereotaxic frame. Craniotomies were made and custom-made tungsten electrode triplets or duplets (0.002 mm diam., California Fine Wire Co.) were implanted into the Primary Motor Cortex (M1; AP: +3.0 mm, ML: ±3.0 mm, DV: −1.4 mm), Primary Somatosensory Cortex (S1; AP: +1.0 mm, ML: ±4.0 mm, DV: −1.4 mm), Dorsal Hippocampus (dHC; AP: −4.5 mm, ML: ±3.0 mm, DV: −2.5 mm), and in some animals into the Primary Visual Cortex (V1; AP: −7.5 mm, ML: ±3.0 mm, DV: −1.4 mm) for recording, and either into the Amygdala (Amy; AP: −4.5 mm, ML: −4.6 mm, DV: -7.5 mm) or Ventral Hippocampal Commissure (vHC; AP: −0.9 mm, ML: - 1.5 mm, DV: −1.5 mm) or Dorsal Hippocampus (dHC; AP: −3.0 mm, ML: −3.0 mm, DV: −3.0 mm) for kindling stimulation; all electrodes were secured with dental cement. Five transcranial electrode screws (Kofuseibyo Co., LTD.) were placed on the temporal/parietal bones bilaterally. After the connectors were soldered and the copper mesh was secured, the animals received antibiotic and analgesic treatment and were housed individually.

#### Kindling procedure

After the 1-week post-surgery recovery period, the animals were connected to the recording system (KJE-1001, Amplipex Ltd, Szeged, Hungary) with a preamplifier headstage and the kindling process started. The dorsal hippocampus was unilaterally stimulated six times daily at 30-min interstimulus intervals with an external stimulus generator (Multi Channel Systems). Each kindling stimulation consisted of 120 biphasic rectangular pulses (0.5 ms positive and 0.5 ms negative) at 66 Hz. The starting amplitude (usually 20–60 μA) depended on the threshold stimulation, where the aim was to evoke trains of after-discharges in most of the hippocampal channels. The amplitude was gently increased over days to a maximum of 40 μA/day. Animals that did not show adequate seizure development were excluded. During and after each stimulation the behavior of the animals was carefully monitored, and seizures were graded from 0 to 5 on the Racine scale^29^. Once the animals experienced at least three generalized motor seizures, they were considered fully kindled. Of the 21 operated animals, eight became reliably fully kindled and yielded electrophysiological recordings of sufficient quality for phase detection and stimulation– response analysis; only these animals were included in the final dataset.

#### Electrophysiological recordings and transcranial stimulations

During the experiments, the animals were kept in their home cages. LFP data were recorded from 32 channels at a sampling rate of 500 Hz (NeuroClinical Desktop Rat Neurostimulator OK-1, Neunos ZRt, Szeged, Hungary) and the behavior was monitored.

After animals reached a fully kindled state, control recordings were obtained from electrically evoked seizures, with approximately 5–10 seizures recorded per animal. These recordings were used both to identify the hippocampal channel exhibiting the most robust and reliable seizure-related activity for phase detection and to manually calibrate the phase detection algorithm on an animal-specific basis, ensuring that phase estimation was tailored to the individual oscillatory characteristics of each subject. The performance of the phase detector was then verified offline to confirm reliable phase tracking before initiating stimulation experiments.

The custom-made phase detector derives the instantaneous phase of the EEG signal on a selected channel by forming an analytic-signal representation via a Hilbert transform. Stimulation is then triggered when the phase reaches a predefined target (e.g., the peak), enabling precise phase-locked timing of the stimulus relative to the ongoing seizure oscillation.

Closed-loop stimulation was administered over a one-week period, with six stimulation trials performed per day. Target phases were scheduled in pseudorandomized order across sessions. Individual trials were excluded before analysis because of damaged or noisy recordings, missing triggers, failed seizure induction, stimulus misalignment, or exploratory stimulation; the resulting session counts are provided in Table S1. Seizure and trial annotations were performed without blinding to stimulation condition or target phase. Given the pronounced behavioral component of the seizures, electrophysiological recordings were occasionally compromised by movement-related artifacts. To minimize animal burden, the number of seizure-evoking stimulations was therefore limited to this maximum of six per day, which in some cases resulted in a reduced number of seizures available for subsequent analysis.

The stimulation parameters were based on those previously used in human experiments ^19^. A half-sine waveform with an amplitude of 5–7 mA was distributed across the stimulation electrodes using the ISP method. This waveform was delivered three times at 95-ms intervals.

#### Histology

Under 2% isoflurane anesthesia, electrode sites were electrically lesioned using 100 μA for 10 s, after which the animals were perfused. The brains were postfixed in 4% formaldehyde, and 50-μm-thick coronal sections were cut at the level of the electrode sites. Sections were stained with 1 μg/ml DAPI and coverslipped, and electrode locations were verified using a Zeiss LSM880 scanning confocal microscope (Carl Zeiss).

### Data analysis

#### Phase detection

For analysis, the exact phase of the seizure oscillation at the seizure onset zone during stimulus delivery was determined offline. First, the phase delay introduced by the recording device filters was corrected using inverse filtering. Then, to reduce noise and slow DC drifts, a bidirectional Butterworth bandpass filter was applied between 1 and 45 Hz. During the analysis, the exact phase of the LFP oscillation targeted by the stimulation was determined as described in the human analysis. Phase analysis was based on a deep hippocampal wire-electrode signal; therefore, all phase information reported below represents oscillations at the seizure-onset zone.

#### Annotation

Seizure segments of the recorded LFPs were annotated manually. Seizure onset was marked at the earliest moment after seizure triggering, when the seizure oscillations appeared at least on one of the channels. Seizure end was marked at the earliest point at which oscillations were no longer present on any channel and ictal behavior had ceased, as confirmed by concurrent video. Generalized seizures were further divided into an initial focal segment, the generalized segment and, when present, a subsequent focal segment.

### Human experiments

#### Human study protocol

The present work is a retrospective, secondary analysis of de-identified recordings acquired within a prospective, single-arm first-in-patient study, registered at ClinicalTrials.gov (identifier NCT07041619). All experimental procedures of the parent study involving human participants were conducted in accordance with the Declaration of Helsinki and approved by the National Institute of Pharmacy and Nutrition and the Medical Research Council of Hungary (Permission Numbers: OGYÉI/9674/2021 and IV/1920-1/2021/EKU). Written informed consent was obtained from all participants following detailed verbal and written explanations of the study’s aims, methodology, potential risks, and anticipated benefits. Participants retained the right to withdraw from the study at any time without consequence. Data collection and all clinical evaluations were conducted at the National Institute of Mental Health, Neurology and Neurosurgery in Budapest, Hungary, now operating as the Clinic for Neurosurgery and Neurointervention, Semmelweis University.

Detailed patient selection, surgery and study design have been described previously by Chadaide et al., 2026^19^. Briefly, participants with clear ictal EEG features and intermittent focal seizures were selected, and potential seizure-onset zones were determined from previous EEG recordings and clinical features. Four eight-contact subgaleal electrode-array strips (MS08R-IP10X-0JH and MS08R298 IP10X-000,

Ad-Tech Medical Instrument Corporation, WI, USA) were implanted subcutaneously on the skull surface as recording and stimulation electrodes. After the baseline recordings, the patients received closed-loop transcranial ISP stimulations with the Neunos SeizureStop device, with concurrent video-EEG monitoring (Neunos ZRt, Szeged, Hungary).

#### Dataset and data analysis

In the human study, participants with epilepsy received ISP stimulation during epileptiform events. Stimulation timing was not immediate upon seizure onset due to the latency of the seizure detection system and the requirement for manual authorization of stimulus delivery by qualified clinical personnel. Consequently, stimulation timing varied considerably following seizure onset. The ensuing analysis examined whether stimulation delivered at different stages of seizures within the seizure-onset zone differentially affected seizure duration. Surface EEG montages substantially influence the vectorial projection of the deep brain dipole onto the recorded surface signals, and may also introduce spatial aliasing manifested as arbitrary phase shifts in the recorded surface projections of the seizure oscillations. These effects can vary across patients and between different seizure sources. To address this uncertainty, we analyzed oscillations reconstructed at the deep-brain source instead of the surface signal. Given the absence of intracerebral signal recordings, the ground truth activity originating from the seizure onset zone was unavailable. Therefore, putative deep-brain activity was reconstructed using the Seizure Phase-transfer Model from the available subgaleal EEG signals^30^ and was used for further analysis (see Figs. 4 and S2).

At the time of this retrospective analysis, five participants from the related first-in-patient study had available, sufficiently complete recordings for seizure-level phase analysis. No additional clinical inclusion criteria were applied beyond eligibility for the parent study; inclusion in the present analysis was determined by data availability and technical suitability for seizure annotation, source reconstruction and phase estimation. Each participant was recorded for 7–8 consecutive days, using the 32-channel EEG system described in Chadaide et al., 2026^19^. Data were collected using four implanted and externalized 8-contact electrode strips (total 32 channels) with a custom-made tabletop EEG recording and closed-loop stimulation delivery device^19^. The EEG signal was recorded at a sampling rate of 500 Hz. Trained epileptologists annotated the recordings, identifying four to 52 seizures per participant. A subset of these recorded seizures received ISP stimulation and was included in the phase analysis.

For each ISP stimulation train, the onset of the first stimulus was defined as the stimulation timing. For seizures where multiple stimulations occurred within a short timeframe (<5 s), only the initial stimulation’s phase was considered for analysis.

Human phase–response relationships were quantified in parallel using two signal representations: the directly recorded surface EEG channel closest to the clinically identified seizure focus and the reconstructed source signal attributed to the seizure-onset zone. The source-reconstructed analysis was used for the primary cross-scale comparison because it provides the closest available approximation to the local population activity represented by the simulated signal and the hippocampal LFP used in rats. The surface-channel analysis was retained as an observation-level robustness analysis and to determine whether phase-dependent susceptibility remained detectable from a directly measurable signal without source reconstruction.

To ascertain the phase of the seizure oscillation at the initiation of stimulation, the surface EEG signal was initially transformed to the deep brain dipole source using a combination of the Gábor–Nelson method and Principal Component Analysis^30^. Subsequently, the reconstructed activity from the seizure-onset zone underwent a phase-correction procedure to mitigate the phase-shift introduced by the analog electronics of the recording device. The phase-shift characteristics of the recording hardware were empirically determined through cadaver head experiments^30^. Following this, the signal was bandpass filtered using a bidirectional Butterworth filter with a frequency range of 1 to 20 Hz.

The final step in quantifying the phase of the surface EEG and the corresponding reconstructed deep-brain activity during stimulation involved the identification of peaks and troughs preceding the stimulation period, their extrapolation into the stimulation period, and interpolation between them at the precise moment of stimulation onset. Peaks and troughs of seizure-related oscillations were detected within a brief time window (approximately 0.5–1.5 seconds) preceding the initiation of ISP stimulation. This window was carefully selected to avoid contamination by stimulation artifacts. Peak detection was performed using the SciPy signal-processing library, with manual parameter tuning to optimize detection accuracy. Following each identified peak, the first subsequent negative deflection was designated as a trough. To characterize the temporal dynamics of the oscillation, peak-to-peak intervals were used to estimate the principal ictal frequency. The positions of the corresponding peaks and troughs during the stimulation period were extrapolated from the final two detected peaks in the pre-stimulation segment. To estimate the instantaneous phase of the oscillations, we employed a Peak-to-Trough-to-Peak (P2T2P) interpolation-based method. Detected peaks were assigned a phase value of 90°, while troughs were assigned 270°, serving as phase anchors. Phase interpolation was performed separately between the peak-to-trough (90° to 270°) and trough-to-peak (270° to 90°) intervals, allowing for asymmetric phase progression across the oscillatory cycle. This approach captures the natural asymmetry often observed in seizure oscillations, where the peak-to-trough interval is typically shorter than the trough-to-peak segment, leading to a non-uniform phase velocity across the cycle.

## Results

### Simulations

Simulations revealed a sharp phase dependence of closed-loop ISP efficacy. In a random network of 100 extended Morris–Lecar populations that produced seizure-like oscillations, we compared matched seizures with and without a single train of phase-locked stimulation (Fig. 1A and B). Of 100 runs, 87 yielded reliable seizures and entered the phase-testing protocol. While the state-space analysis is illustrated using an example seizure with stimulations applied at 60 different phases (Fig. 1C–G), the systematic evaluation of seizure-length dependence on stimulation phase was performed by selecting 8 stimulation phases for each of the 87 seizures. These phases were sampled uniformly by drawing one phase from each of eight 45° bins per simulation, yielding a total of 696 stimulated trials. Seizure durations were normalized to the median duration of the 87 unstimulated seizures.

**Figure 1.**
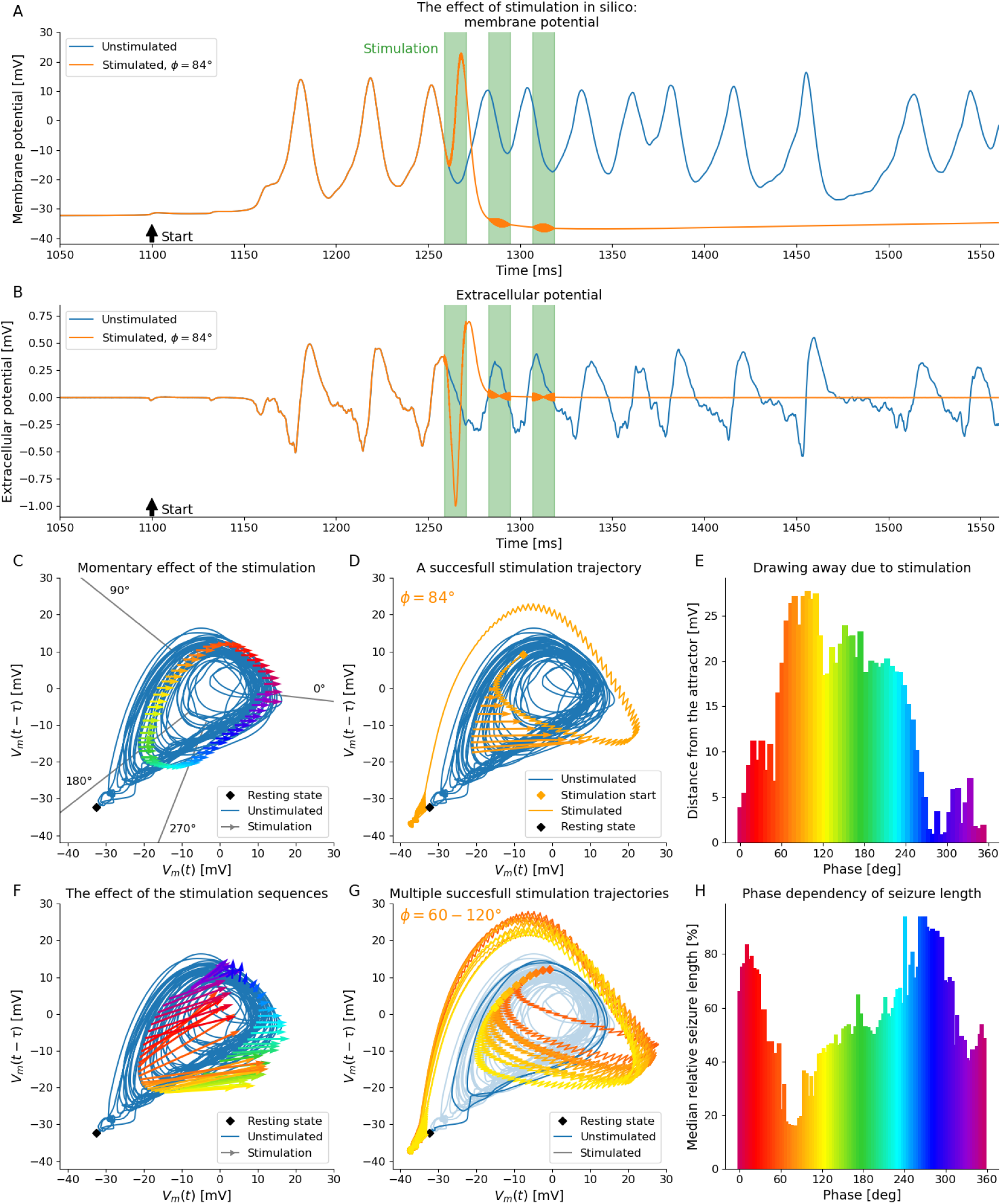
Effect of stimulation on a simulated seizure. **(A)** Example time series of mean membrane potential of the modeled neural population. Comparison of the unstimulated (blue) and stimulated seizures (orange) during simulation. The time of stimulation is denoted by green stripes**. (B)** The parallel simulated extracellular potential. **(C)** State-space reconstruction of the epileptic dynamics through time-delayed embedding of the mean membrane potential (blue whirlpool). Colored arrows show the momentary effect of stimulation initiated at different phases. **(D)** The temporal evolution of a prolonged stimulation effect (orange arrows) led the system to a new trajectory (orange line) on which the seizure stops and the system returns to the resting state within one period. **(E)** Displacement from the original seizure attractor as a function of the stimulation starting phase. **(F)** Phase dependence of the prolonged stimulation effect. **(G)** Multiple successful stimulated trajectories. **(H)** Median seizure lengths in a 50° wide sliding window as a function of stimulation starting phase. The most effective stimulation trains started between 60° and 90°, where the displacement (on graph E) was maximal.

Matched pairs confirmed that a single phase-locked train can terminate a seizure within one cycle whereas the unstimulated counterpart persists. In representative trials, the stimulated and control trajectories were identical up to stimulation onset and then diverged abruptly, with rapid seizure termination in the stimulated run (Fig. 1A–B). These examples illustrate the magnitude and speed of the state change the control rule is designed to exploit.

State-space analysis linked efficacy to how ISP displaces the ictal trajectory. Time-delay embedding^25^ revealed a low-dimensional limit-cycle-like attractor for the seizure dynamics. Brief pulses produced an instantaneous rightward shift (toward higher V) that was largely independent of the starting phase (Fig. 1C). The cumulative displacement caused by a stimulus train pushed the trajectory across the basin boundary into the resting fixed point, yielding termination within a single cycle (Fig. 1D).

Mapping the prolonged effects across various target starting phases showed that trajectories initiated around the rising/peak segment escaped the attractor most often, whereas stimulations near the trough had little effect. Aggregated trajectories colored by starting phase emphasize successful exits for ∼60–120° starts and minimal impact for ∼270–360°. Early-rising starts (∼0–60°) produced large displacements that nevertheless fell short of the basin boundary (Fig. 1F–G). This directional asymmetry explains why only a subset of large momentary displacements translate into termination.

The instantaneous displacement itself was phase-modulated and predicted efficacy. The 10-ms post-onset distance from the unstimulated attractor peaked for stimulations initiated at ∼60–120° (Fig. 1E), mirroring the phase window that most frequently produced escapes in Fig. 1F–G and the shortest seizures in Fig. 1H. Quantitatively, the normalized median seizure duration exhibited a broad optimum around 60–90° with a global minimum at 79.2°, declined across the falling phase, and entered a largely ineffective regime near ∼270°, with a shallow secondary minimum near ∼330° (Figs. 1H, S3 and S4). Together, Fig. 1E–H establish that phase-maximal displacement coincides with phase-maximal shortening.

### Animal experiments

To assess the efficacy of ISP-phase-targeted stimulation in suppressing epileptic seizures, we first compared full and generalized seizure durations between stimulated and non-stimulated control conditions across multiple animals. The stimulation resulted in a marked reduction in full seizure length with statistically significant differences in 7 of the 8 rats (Fig. 2A and B, Table S2; Mean normalized length = 100 ± 56 % for control group, and 69 ± 39 % for stimulated group, N = 167 control vs 104 stimulated seizures in 8 animals, each seizure normalized to the median control length in the given animal; P = 6.63 × 10^−14^, Kolmogorov-Smirnov test). Generalized seizure durations showed a similar trend with shorter durations in the stimulated condition compared to control, indicating that ISP stimulation can attenuate both local seizure expression and broader propagation (Fig. 2C and D, Table S3; Mean normalized length = 100 ± 36 % for control group, and 75 ± 36 % for stimulated group, N = 167 control vs 104 stimulated seizures in 8 animals, each seizure normalized to the median control length in the given animal; P = 1.28 × 10^−8^, Kolmogorov-Smirnov test).

**Figure 2.**
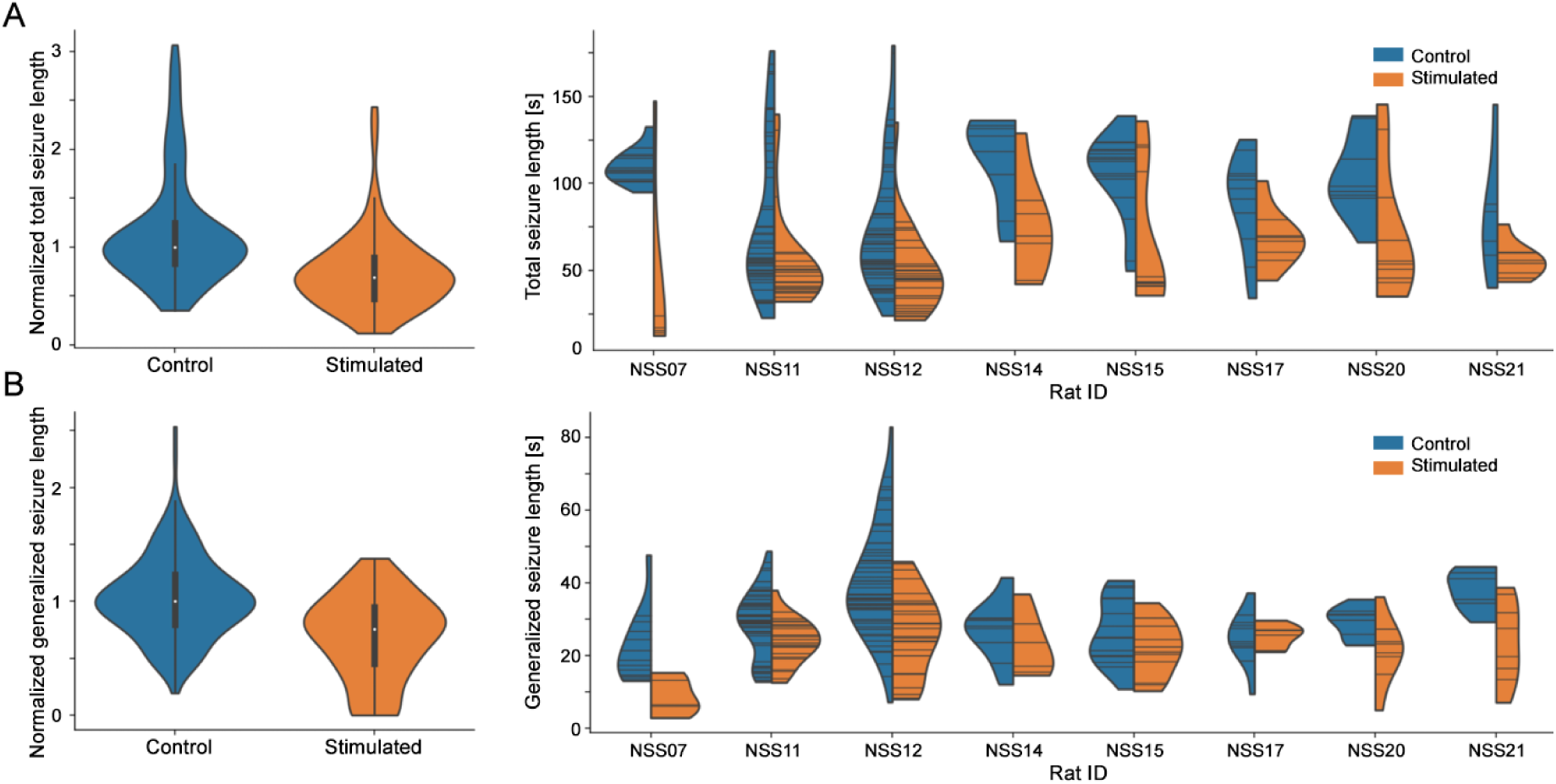
ISP stimulation reduces seizure durations across full and generalized events. **(A)** Violin plots show the distribution of normalized full seizure durations in seconds under control (blue) and ISP-stimulated (yellow) conditions, both pooled (left) and broken down by individual animals (right). **(B)** Generalized seizure durations under the same conditions, again shown pooled (left) and by individual animals (right). In both seizure categories, ISP stimulation resulted in a reduction in seizure length compared to control trials. While individual variability was observed, the general trend across animals consistently showed shorter seizures with ISP stimulation.

We next examined whether the timing of ISP stimulation relative to the ictal phase of seizure activity modulated its effectiveness. Stimulation was most effective when delivered around 45° and 315° of the ictal phase (Fig. 3B and S5; Table S4). This timing corresponded to the rising phase of the seizure waveform, illustrated schematically by the red sinusoidal trace. Median full seizure duration dropped to 57.57% of control at 45° and 56.28% at 315°, with Bonferroni-corrected P = 1.22 × 10^−4^ and P = 3.10 × 10^−5^. Other phases, including 90° and 135°, also produced significant reductions compared to control. Durations of the generalized periods followed a similar but attenuated trend (Fig. 3C and S6; Table S4). The strongest effect again occurred at 45°, with median generalized seizure length reduced to 41.69% (Bonferroni-corrected P = 3.71 × 10^−4^).

**Figure 3.**
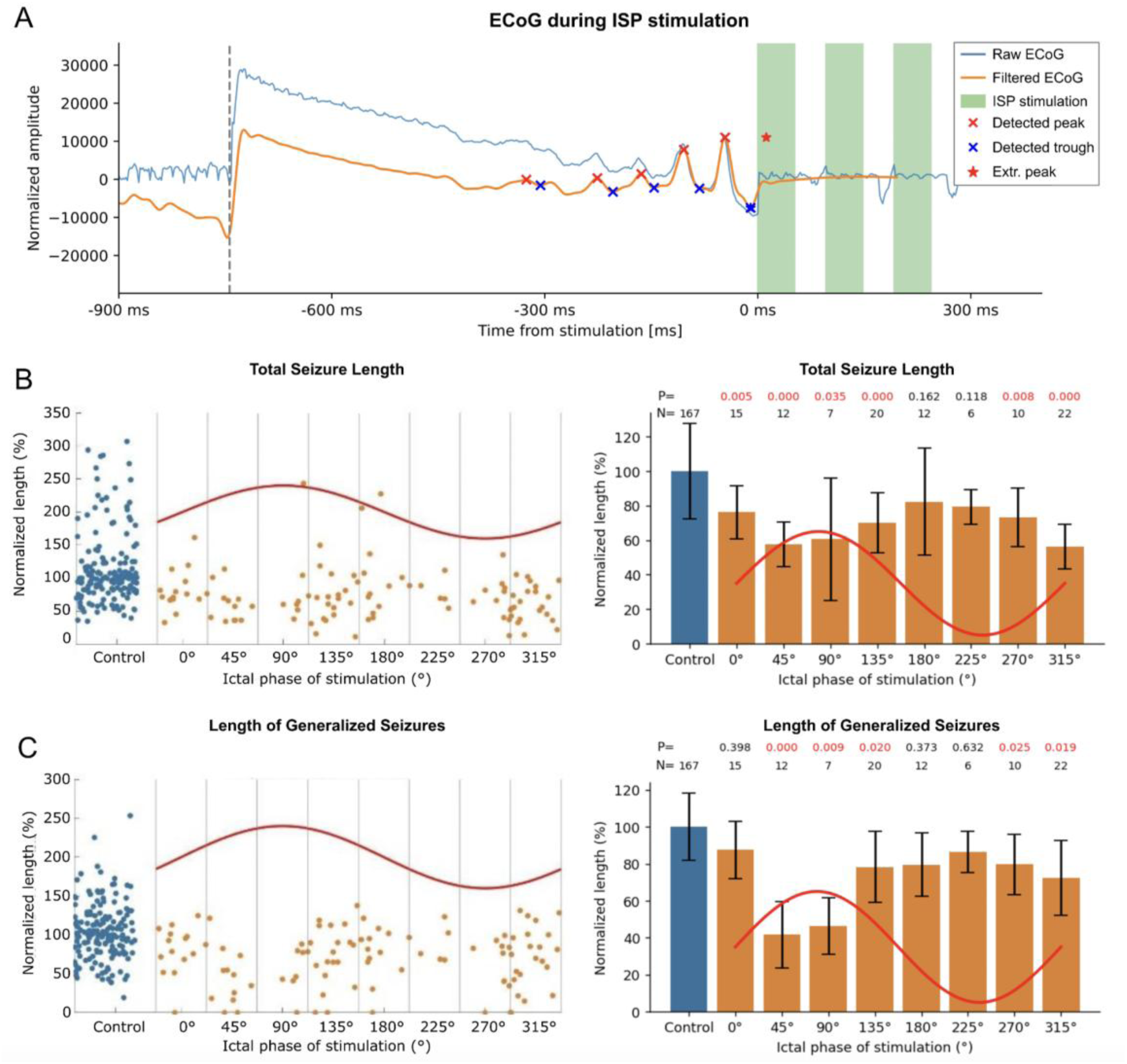
Seizure suppression depends on the ictal phase of ISP stimulation in rats. **(A)** A sample illustrating phase estimation. After phase correction and filtering of the original signal (blue and orange traces, respectively), peaks and troughs (red and blue markers) were detected. These are extrapolated beyond the start of the stimulation (red star during the stimulation, which is marked by green bars) and the phase values between the last detected (’x’) and the extrapolated peak (asterisk) are interpolated. **(B)** Normalized full seizure lengths under ISP stimulation delivered at different ictal phases (in 45° increments, yellow), compared to control (no stimulation, blue), show consistent phase-dependent effects, with the strongest seizure suppression occurring when stimulation was applied near 45°. **(C)** Generalized seizure durations similarly depended on stimulation phase, with largest seizure shortening when stimulating at 45°. For B and C, left panels depict single-trial data, while the right panels show mean ± SEM for each phase. The red sinusoidal curve schematically represents the underlying oscillatory activity of the seizure, illustrating stimulation timing relative to ictal phase.

**Figure 4.**
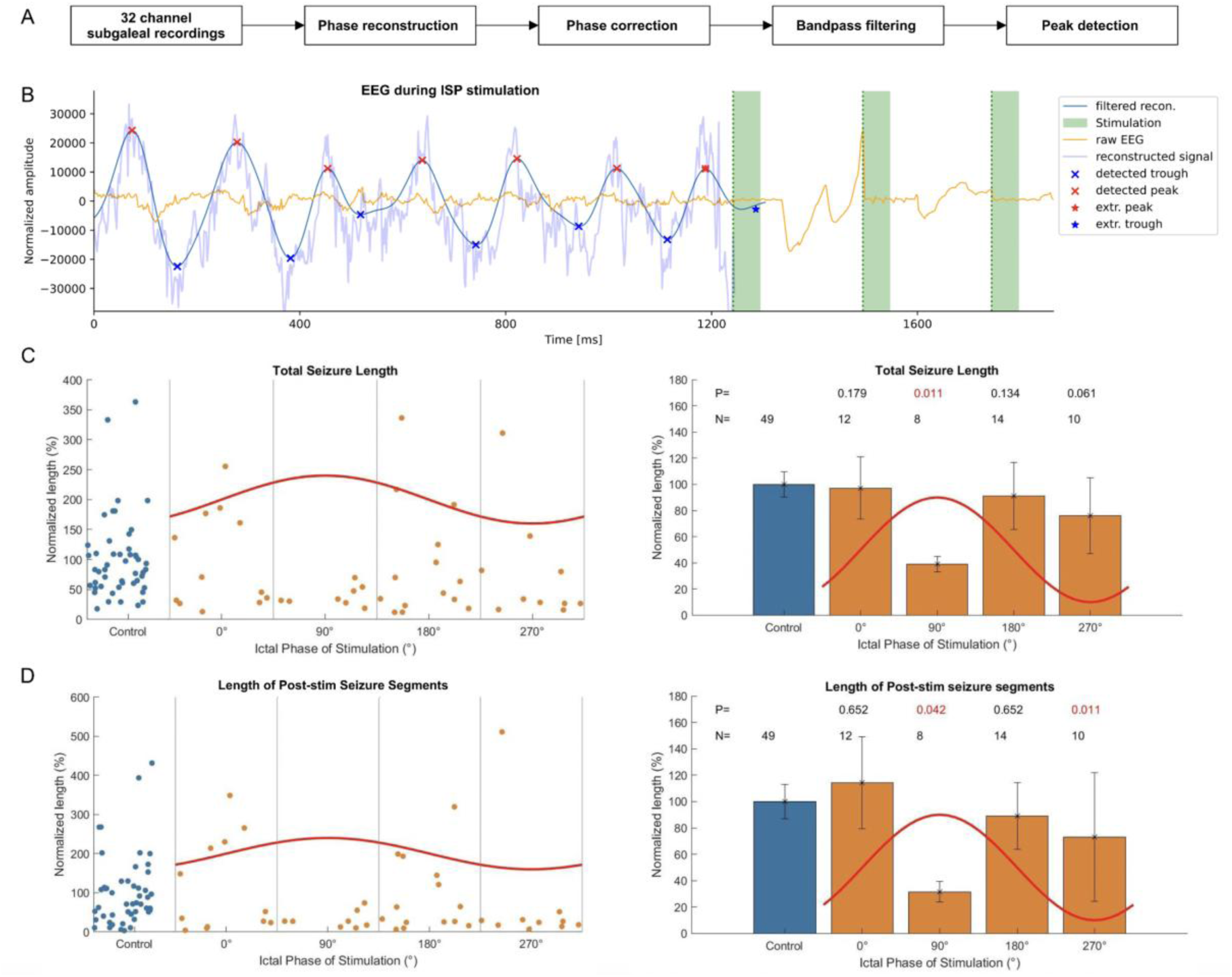
Seizure suppression depends on the ictal phase of ISP stimulation in humans. **(A)** Data processing pipeline. **(B)** Phase estimation example: the raw signal from a subgaleal-EEG channel (orange) of Patient #3 is shown together with the source signal reconstruction (light blue) generated using the Seizure Phase-transfer Model. Following phase correction and band-pass filtering (dark blue), peaks (red markers) and troughs (blue markers) are identified. The peaks are extrapolated beyond the start of stimulation (green columns), and a phase value is subsequently estimated. **(C, D)** Change in total seizure duration (C) and post-stimulus seizure duration (D) as a function of the targeted ictal phase of the reconstructed deep-brain dipole. Data are normalized to each patient’s average control seizure length. Right panels show the quantized measures (mean ± SEM) of the individual seizure data shown on the left panels as scatter plots. Unstimulated control cases are shown in blue.

Notably, the magnitude of reduction in generalized segments was generally smaller and more variable than in full seizures. Thus, the rodent phase-response profile was non-uniform rather than confined to one narrow phase angle: the strongest effects occurred at 45° and 315°, with additional significant shortening at neighboring phases.

Together, these findings demonstrate that ISP stimulation reliably shortens both the overall duration of the seizures and the lengths of the generalized periods. Its efficacy is strongly influenced by the timing of delivery relative to the phase of ongoing seizure activity. Early-phase stimulation - particularly near 45° - is associated with the most potent seizure-suppressing effects.

### Human experiments

We first examined the association between ISP-induced seizure shortening and the ictal phase targeted by stimulation by analyzing stimulation timing relative to the reconstructed activity of the seizure-onset source, which provided the signal representation most directly comparable to the local population signals used in the simulations and rat experiments (Fig. 4A, B). Given the limited number of total seizures, data were aggregated across all patients. Prior to aggregation, seizure lengths were standardized by the mean duration of each patient’s unstimulated control seizures.

To investigate the phase-specific efficacy of seizure interruption, we compared the full stimulated seizure durations across four stimulation phase bins, centered at 0°, 90°, 180°, 270°, relative to a non-stimulated control condition. Mean seizure length in the control condition was standardized to 100% (SEM = 9.82%, N = 49). Stimulation around 90° - corresponding to the peak of the oscillation - resulted in the most substantial reduction in seizure duration, with a mean length of 39.14% (SEM = 5.89%, N = 8) (Fig. 4C). This effect was statistically significant when compared to control (P = 0.0028), and remained significant after Bonferroni correction (P = 0.0113). Stimulation at 270° (trough) was associated with a reduction in seizure duration (mean = 75.90%, SEM = 28.93%, N = 10; uncorrected P = 0.0204), although this effect did not remain significant after Bonferroni correction (corrected P = 0.0613). In contrast, stimulation at 0° (mean = 97.25%, SEM = 23.55%, N = 12) and 180° (mean = 91.03%, SEM = 25.64%, N = 14) produced minimal effects on seizure length, with no statistically significant differences from the control group (Table S5). These results support a phase-dependent effect of targeted ISP stimulation, with the greatest seizure suppression observed when stimulation was precisely timed around the peak of the ictal oscillation. The phase effect was also examined using pairwise bin comparisons (Fig. S7).

To account for the variability in stimulation delivery delays occurring after seizure onset, we also analyzed seizure durations following the initial stimulus (Fig. 4D and S8). For this analysis, the control seizures’ durations were reduced by the average stimulus delivery delay of the given patient prior to individual normalization. The control condition (no stimulation) was standardized to 100% (SEM = 13.17%, N = 49). Stimulation at 90° (peak phase) again resulted in the most pronounced reduction in seizure duration, with a mean remaining seizure length of 31.57% (SEM = 7.80%, N = 8), representing a statistically significant reduction relative to control (P = 0.0138; Bonferroni-corrected P = 0.0415). Stimulation at 270° (trough phase) similarly resulted in a significant reduction (mean = 73.10%, SEM = 48.79%, N = 10; P = 0.0028; Bonferroni-corrected P = 0.0113). In contrast, stimulation at 0° (mean = 114.26%, SEM = 34.96%, N = 12) and 180° (mean = 89.15%, SEM = 25.14%, N = 14) did not produce significant differences in seizure duration compared to control (Table S6). These findings reinforce the phase-specific modulation of seizure termination, with stimulation delivered at peak or trough phases yielding the greatest reductions in ongoing seizure activity. The human data therefore identify 90° as the most prominent efficacy bin, while also indicating a possible second susceptible region around the trough; this latter effect should be interpreted cautiously given the smaller sample sizes and variability within phase bins.

Consistently, both the overall seizure duration analysis and the post-stimulation seizure duration analysis demonstrated that targeting the ictal oscillations of the seizure onset zone at or slightly preceding their peak phase is more effective in shortening or terminating seizures than stimulation delivered at other phases.

As a complementary observation-level analysis, we repeated the complete phase–response analysis using the directly recorded surface EEG channel showing the clearest seizure oscillation (Fig. S9). This analysis also revealed phase-selective seizure shortening, but the apparent efficacy map was displaced relative to that obtained from the reconstructed seizure-onset-zone source. Total seizure duration was reduced to 48.25 ± 11.91% of control when stimulation was delivered at 0°, corresponding to the rising zero-crossing of the surface oscillation (N = 11; Bonferroni-corrected P = 0.0314), and to 69.28 ± 21.24% at 270°, corresponding to the trough (N = 15; corrected P = 0.0027). In contrast, stimulation at the surface-defined peak (90°; 105.54 ± 20.15%, N = 9; corrected P = 0.7092) or at 180° (110.42 ± 41.22%, N = 9; corrected P = 0.0535) did not significantly shorten seizures relative to the unstimulated control condition.

The analysis of seizure duration remaining after the first stimulation yielded the same surface-phase pattern (Fig S10). Remaining duration was reduced to 39.22 ± 12.22% of control at 0° (N = 11; corrected P = 0.0393) and to 78.72 ± 34.68% at 270° (N = 15; corrected P = 0.0121), whereas no significant reduction was observed at 90° (122.71 ± 38.34%, N = 9; corrected P = 0.6536) or 180° (98.46 ± 40.13%, N = 9; corrected P = 0.4876).

Thus, stimulation efficacy remained related to the phase measured directly at the surface, but the angular location of the effective windows differed from the peak-centred susceptibility identified in the reconstructed source signal. Complete surface-channel distributions, pairwise comparisons and descriptive statistics are provided in Figs. S9 & S10 and Tables S7 & S8.

## Discussion

The present study demonstrates that ISP stimulation can shorten epileptic seizures in a phase-dependent manner across computational models, rodent experiments, and human recordings. By systematically comparing stimulation at different oscillatory phases, we found strongly non-uniform phase-response profiles in all three preparations. A prominent efficacy region appeared in the rising-to-peak portion of the seizure cycle, but the exact optimal angle and the presence of secondary susceptible regions differed between simulations, rodents and humans. These findings extend prior demonstrations of closed-loop stimulation by adding an additional layer of temporal precision, supporting the concept of phase-targeted neuromodulation.

Stimulation may be particularly effective during the rising and peak phases because these can coincide with heightened network excitability and dynamical instability. However, the secondary susceptible regions observed near the trough in some analyses indicate that susceptibility cannot be inferred from phase alone. At 45°, the strongest relative reduction was observed for generalized seizure segments, suggesting that timely intervention at these phases may be especially effective at disrupting large-scale synchronization. This aligns with recent theories of seizure propagation, which propose that generalized spread depends on the reinforcement of oscillatory synchrony; by perturbing at the critical buildup phase, ISP may weaken the recruitment of additional networks^18,31^.

In the simulations, time-delay embedding revealed a low-dimensional ictal attractor and made the control problem concrete: ISP acts as a directional “kick” that must push the trajectory across the basin boundary. Brief pulses shift the state rightward largely independent of starting phase, but trains initiated on the rising/peak segment accumulate displacement and consistently escape to the resting fixed point. The displacement peaks at ∼60–120° and predicts seizure shortening, implying a minimal control-energy region in that window. The state-space trajectories highlight that a temporary increase in oscillation amplitude can precede termination when stimulation begins in the optimal window. In the model, this overshoot does not indicate a proconvulsant effect; rather, it reflects a directed push across the separatrix into the resting basin. This state-space view may serve as a mechanistic design principle for tailoring stimulation policies.

Rinzel (1985) demonstrated the existence of a bistable regime in the Morris–Lecar model and subsequently in other neural models, in which a stable fixed-point attractor coexists with a stable periodic firing (oscillatory) attractor^32^. His simulations further showed that a single pulse of stimulation can switch the neuron’s behavior: a pulse can push the system from the resting state into the periodic oscillatory regime, where it remains without any additional input, while another pulse can return the system to the resting state, where it also persists.

The imperfect alignment of exact phase optima across simulations, rodents and humans is expected. Phase was estimated from different observables in systems with distinct network architectures, frequency evolution and measurement constraints; the same nominal phase angle therefore need not correspond to an identical latent network state. The translational implication is consequently not that one universal angular target should be applied across all seizures, but that phase-resolved stimulation can identify periods of increased susceptibility within a given preparation or patient. The recurrent prominence of the rising-to-peak region nevertheless provides an empirically useful starting window for real-time targeting.

A closely related representation effect was evident within the human dataset itself. Parallel analysis of the directly recorded surface EEG supported an association between stimulation efficacy and oscillatory phase, but revealed a markedly shifted phase–efficacy map. Whereas shortening was greatest around the peak of the reconstructed seizure-onset-zone source, the surface-channel analysis identified significant effects at the rising zero-crossing and trough, but not at the surface-defined peak. This discrepancy is expected because a surface EEG channel represents a geometry- and reference-dependent projection of the underlying generators: source orientation, volume conduction, electrode montage and spatial aliasing can shift or invert the apparent phase. Moreover, the surface channel with the clearest seizure oscillation was selected individually and, in some instances, differed between seizures. A nominal phase angle measured at the surface should therefore not be interpreted as a fixed angular surrogate of local ictal phase.

The convergence of both representations on phase-selective efficacy supports the underlying biological phenomenon, while their angular displacement shows that its observed phase coordinate depends on how network activity is measured. The reconstructed source estimate is consequently more appropriate for comparison with the local population signals used in the simulations and rat experiments, although it remains model-derived and was not validated against concurrent intracranial recordings in these participants. Translation to prospective surface-EEG control will therefore require either source-aware phase estimation or patient-, focus- and montage-specific calibration of the effective surface phase.

The state dependence of the response of an epileptic system to stimuli was examined by Suffczynski et al. (2004) and Taylor et al. (2014) in computational models of absence epilepsy^8,33^. In this work, we extended the analysis of phase-dependent responses to ISP stimulation by directly comparing model-based predictions with experimental observations obtained from both a rodent epilepsy model and human participants with epilepsy. The convergence of phase dependence across the in silico, rodent and human data suggests that state-dependent stimulation susceptibility is shared across systems, even though its precise phase expression may differ between preparations, and that a deeper understanding of it may lead to more effective seizure- termination strategies.

Because the effective window occupies a fraction of the ictal cycle, end-to-end detection-to-stimulus latency must fit inside that window to hit the target reliably. The manual authorization step in the human study produced variable delays. In deployment, an automated controller should compensate the latency by predicting phase forward and gate delivery only when phase, amplitude, and instantaneous frequency jointly indicate proximity to the basin boundary. This argues for “predict-then-trigger” rather than “detect-and-fire” logic.

Simulations showed that efficacy is not strictly monotonic with amplitude, suggesting iso-efficacy curves over phase and amplitude. Operating near the rising/peak window appears to lower the amplitude needed to cross the basin boundary, offering a practical route to reduce delivered charge and scalp sensation while preserving seizure control. Future protocols should map patient-specific phase– amplitude surfaces and select the lowest-energy setting that achieves escape with high probability. Ideally, a controller should learn a per-patient phase-efficacy function online, through a short calibration phase. The reconstruction of deep-brain phase from subgaleal EEG^30^ may provide the necessary state variable for such adaptive policies.

While our data provide strong support for optimizing ISP stimulation, several limitations need to be acknowledged.

First, the sample size in both rodents and humans was modest, which constrains the generalizability of the findings. Furthermore, our human cohort included different epilepsy types, but the small sample size precluded systematic comparison of whether phase dependence varies across epilepsy types. These limitations highlight the need for future investigations across several animal models (e.g., kainate, pilocarpine, kindling) and diverse human epilepsy types, to establish whether phase dependence represents a generalizable principle or one constrained by pathology and network architecture.

A critical open question concerns the scope of phase-targeted stimulation effects beyond the acute seizure period. While the present analysis demonstrates robust phase-dependent seizure shortening on an acute timescale, it is not designed to assess whether closed-loop ISP intervention influences seizure incidence, interictal activity, or baseline epileptic dynamics over longer durations. Consequently, the current findings should be interpreted as evidence for acute modulation of seizure duration rather than long-term seizure suppression. Whether repeated phase-targeted stimulation produces sustained changes in seizure burden, or whether phase specificity remains relevant outside the immediate seizure context, can only be addressed in future longitudinal and chronic stimulation studies.

Another difference between the experimental settings lies in how many stimulations were delivered per seizure. In the computational models and rodent experiments, ISP was applied only once, isolating the effect of a single phase-targeted train, whereas in humans some seizures received multiple consecutive stimulations. This complicates interpretation, as repeated stimulations may interact or prolong effects, while single applications reveal the immediate impact of phase timing. Future studies should directly compare single versus repeated ISP interventions to test whether benefits accumulate linearly or involve nonlinear interactions.

A feasible method for repeated phase-controlled stimulation is a brief “phase-tiling” sequence that straddles the predicted window (e.g., two or three short trains separated by one-cycle intervals) to hedge against frequency drift and estimator jitter, while keeping total charge within tolerability. The state-space framework predicts diminishing returns once escape is achieved; thus, the algorithm should terminate stimulation immediately after the first successful reset to avoid unnecessary pulses.

Despite these limitations, the present work represents the first systematic investigation of phase-dependent ISP stimulation across modeling, animal experiments, and human data. By demonstrating that the efficacy of ISP depends not only on whether stimulation is delivered in a closed-loop fashion, but also on *when* within the oscillatory cycle it is applied, this study brings a new level of temporal precision to non-invasive neuromodulation. These findings mark an important step toward developing more targeted and effective interventions for people living with drug-resistant epilepsy.

## Acknowledgments

This work was supported by the Momentum program II of the Hungarian Academy of Sciences (AB), EFOP-3.6.1-16- 2016-00008 (AB), EFOP 3.6.6-VEKOP-16-2017-00009 (AB), KKP133871/KKP20 (AB), and 151490/EXCELLENCE_24 (AB) grants of the National Research, Development and Innovation Office, Hungary (AB), the 20391-3/2018/FEKUSTRAT of the Ministry of Human Capacities, Hungary, the EU Horizon 2020 Research and Innovation Program (No. 739593—HCEMM to AB), Ministry of Innovation and Technology of Hungary grant (TKP2021-EGA-28 to AB), Hungarian Scientific Research Fund (grant K135837 to ZS), the Hungarian Brain Research Program (grant NAP2022-I-7/2022 to AB), Hungarian Research Network TECH-2024-020 (ZS).

## Author Contributions

A.B. conceived the project.

L.B., Z.S., Z.Ch. and A.B. developed the experimental protocols and methodology.

L.B., N.F., P.R., and A.B. performed the rodent experiments. Z.S., V.H., and K.F. performed simulation experiments.

L.B., N.F., Z.Ch., T.L., M.H.K., and A.B. analyzed the rodent and human data.

L.B., Z.S., M.H.K., and A.B. wrote the manuscript. L.E., Z.Ch. and A.B. supervised the project.

## Data and Code Availability

The data supporting the findings of this study are available from the corresponding author upon reasonable request. Individual-level human EEG recordings and associated clinical metadata are not publicly available because of participant privacy, consent, and ethical-governance restrictions; access may be considered subject to approval by the relevant data controller and ethics bodies. The simulation and analysis code are available from the corresponding author upon reasonable request.

## Supplementary Material

**Figure S1.**
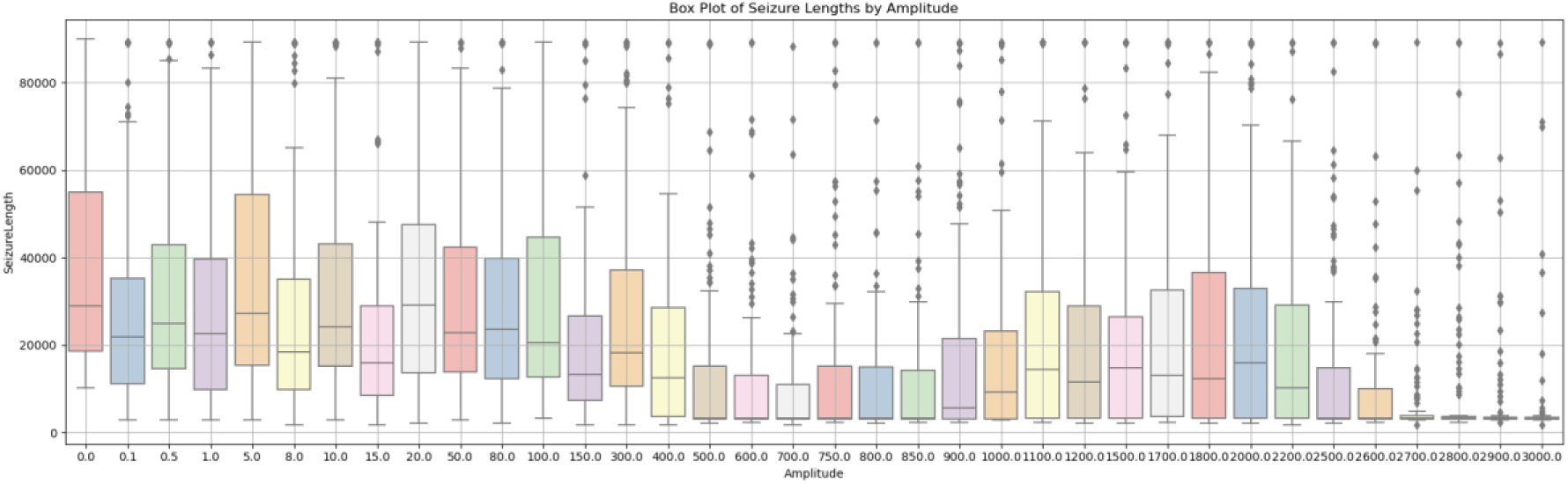
Amplitude dependency of seizure length in simulations. Each bar shows median (black bar), interquartile interval (colored box), 1.5 times the interquartile range (whisker) and the outliers (diamonds). Note, the non-monotonic dependency of seizure-shortening effect on stimulation amplitude. Based on these results, subsequent simulations used a stimulation amplitude of 500.

**Figure S2.**
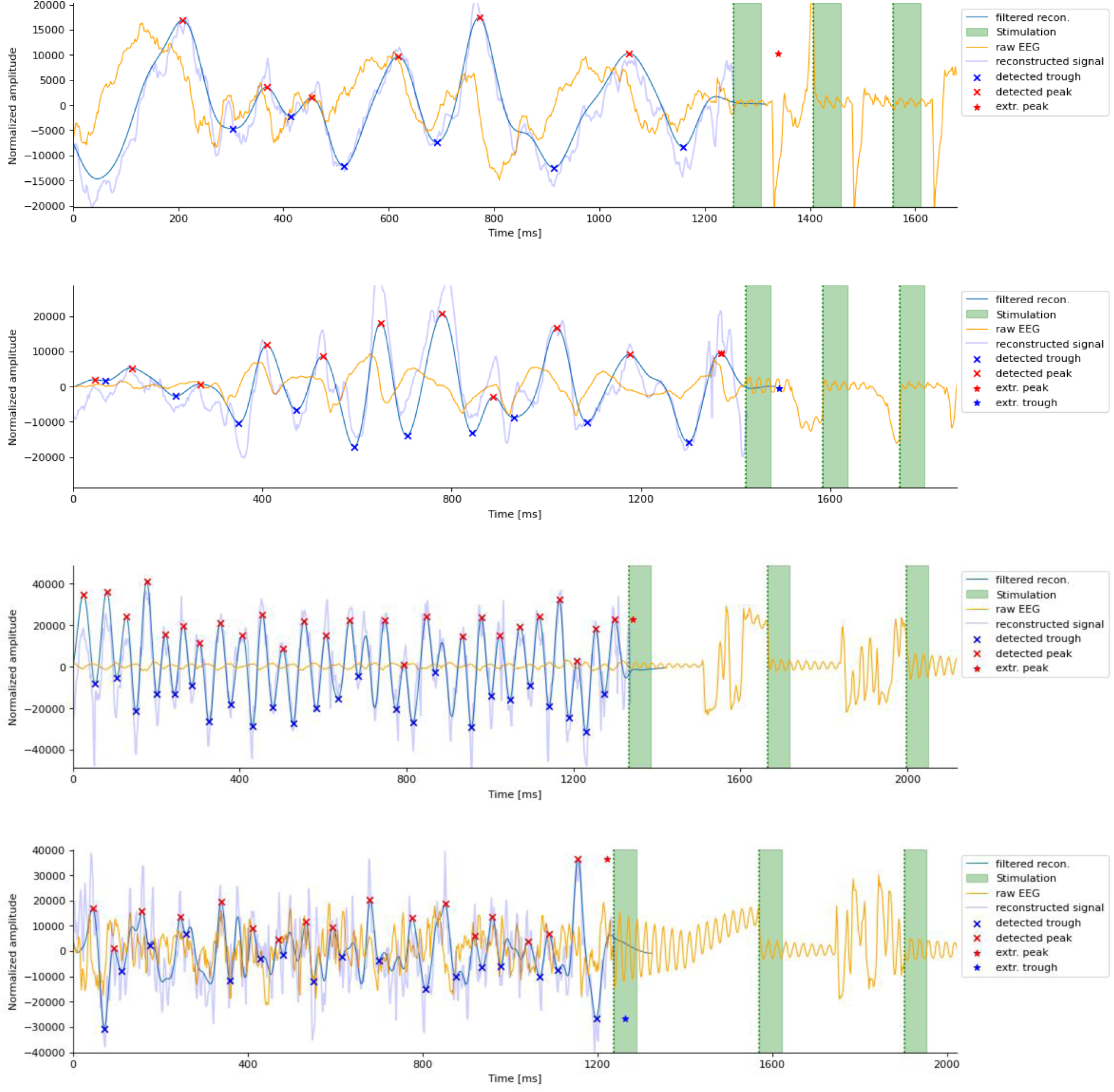
Example segments of source signal reconstruction and phase estimation for each patient. The raw signal from a subgaleal-EEG channel (orange) is shown together with the reconstructed source signal (light blue) generated using the Seizure Phase-transfer Model. Following phase correction and band-pass filtering (dark blue), peaks (red markers) and troughs (blue markers) are identified. The peaks are extrapolated beyond the start of stimulation (green columns), and a phase value is subsequently estimated. From top to bottom: Patient #1, Patient #2, Patient #4, Patient #5 (for Patient #3 see Fig. 4).

**Figure S3.**
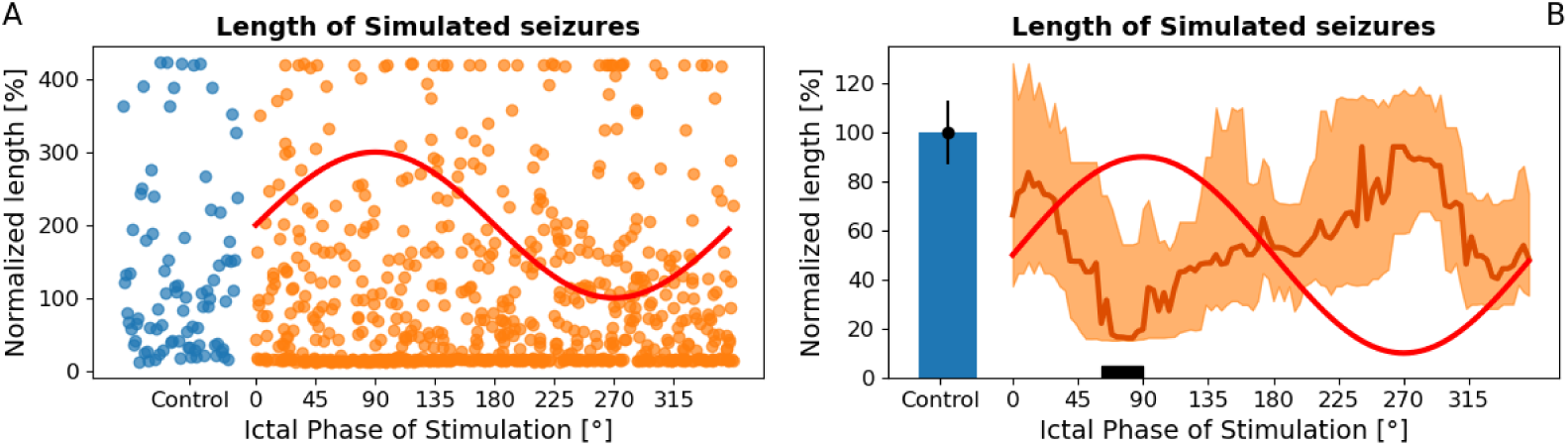
Systematic evaluation of the seizure length dependency on stimulation phase. **(A)** Scatter plot of 87 unstimulated control seizure lengths (blue dots) normalized to their median compared to the stimulated seizure lengths normalized to the median of the control seizures as well. Each seizure was stimulated in 8 different phases that are uniformly distributed within the eight 45°-wide bins, resulting in 696 lengths altogether. Stimulation phases are measured compared to the filtered simulated EC potential, which is schematically marked by the red sine wave. As a result of the stimulation, many seizures remained very short; however, a few long seizures appeared as well, forming a highly skewed seizure length distribution. **(B)**: Median and its 95% confidence interval of the normalized seizure lengths in a 50° wide sliding window, compared to the median and the SE of the unstimulated control seizure length. The seizure length phase dependency shows multiple features similar to the experimental results: stimulation significantly shortened the seizure lengths in most cases and this effect shows strong dependence on the phase of the stimulation: the median seizure length shows two minima: a deeper, optimal stimulation period between 60° and 90° marked by a black rectangle and with the global optimum at 79.2°, and a shallower local optimum between 310° and 360°.

**Figure S4.**
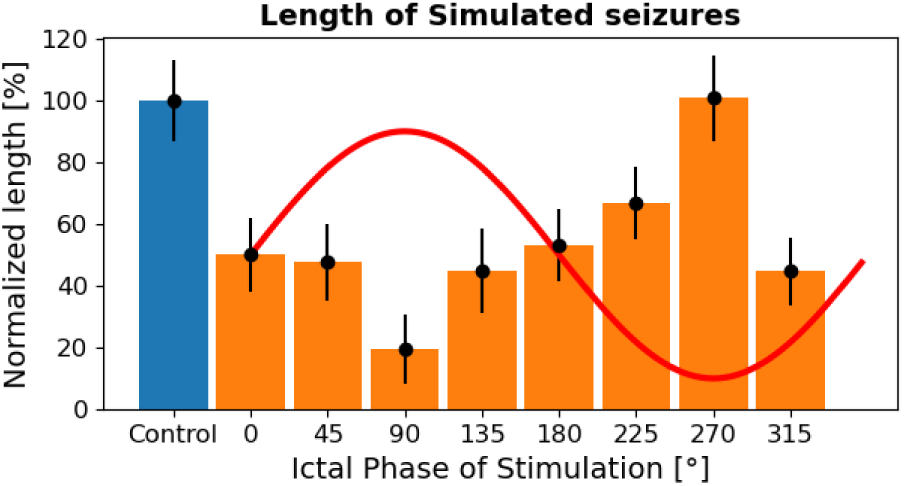
Median of normalized seizure lengths and standard errors within eight 45°-wide bins obtained from the simulations. This representation is directly comparable and shows clear similarity to the experimental results on rats.

**Figure S5.**
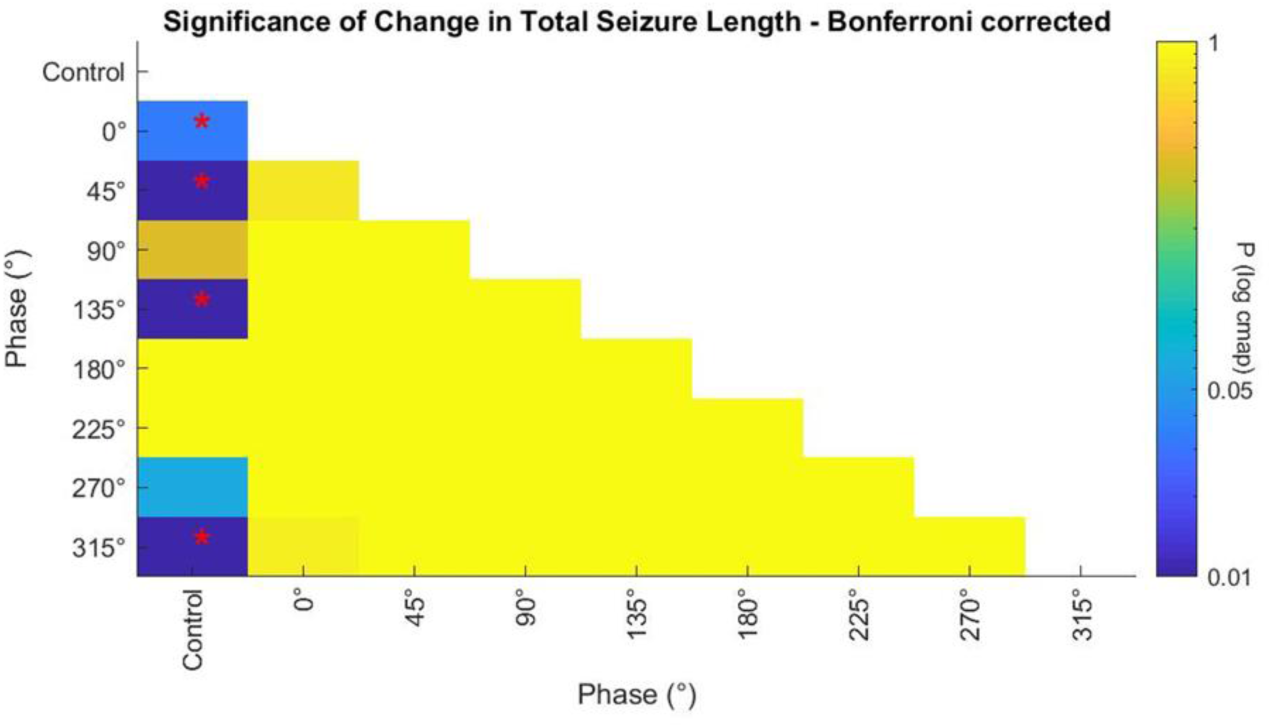
Total seizure lengths in rats - Statistical comparison of efficacy in various phase bins and control regarding the total seizure durations in rats. The P-values of the Bonferroni-corrected pairwise comparisons of total seizure lengths are displayed on a logarithmic color scale for better visibility. Significant pairs (i.e. P < 0.05) are also marked with red asterisks for better readability.

**Figure S6.**
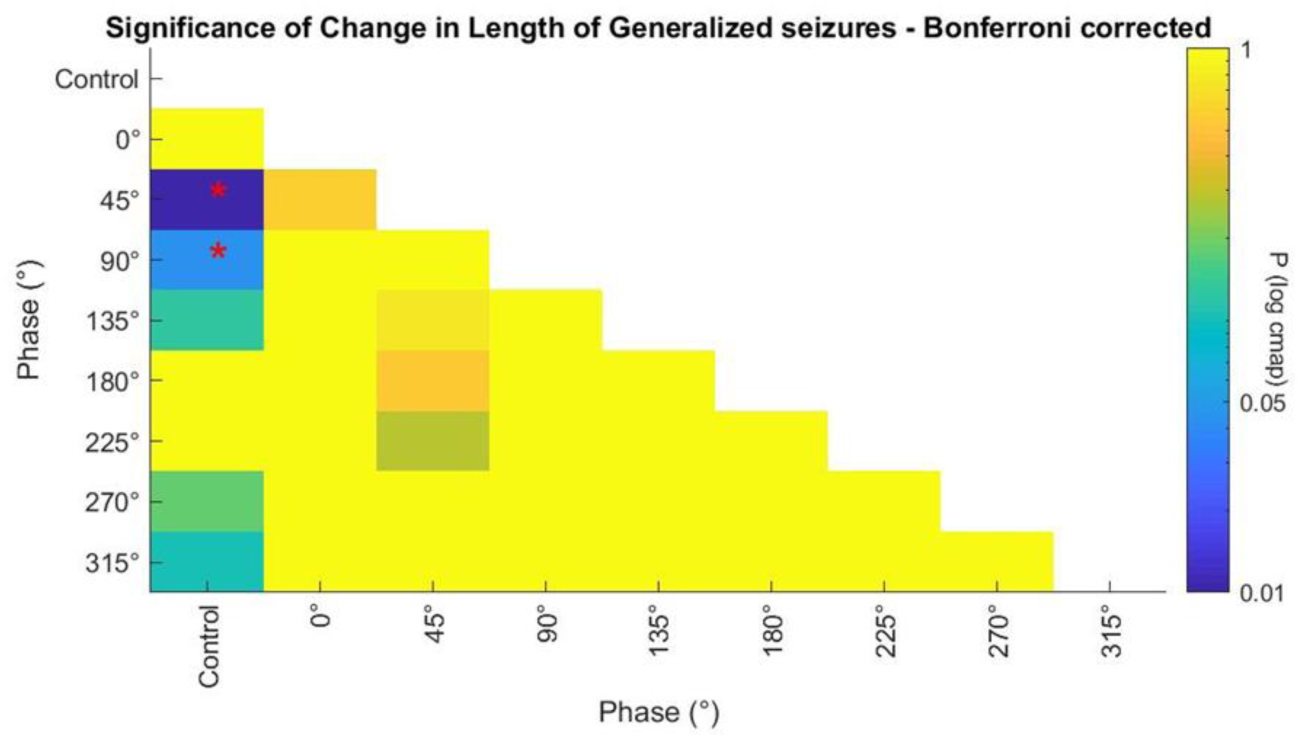
Statistical comparison of efficacy in various phase bins and control regarding the duration of generalized seizures segments in rats. The P-values of the Bonferroni-corrected pairwise comparisons of the lengths of generalized segments are displayed on a logarithmic color scale for better visibility. Significant pairs (i.e. P < 0.05) are also marked with red asterisks for better readability.

**Figure S7.**
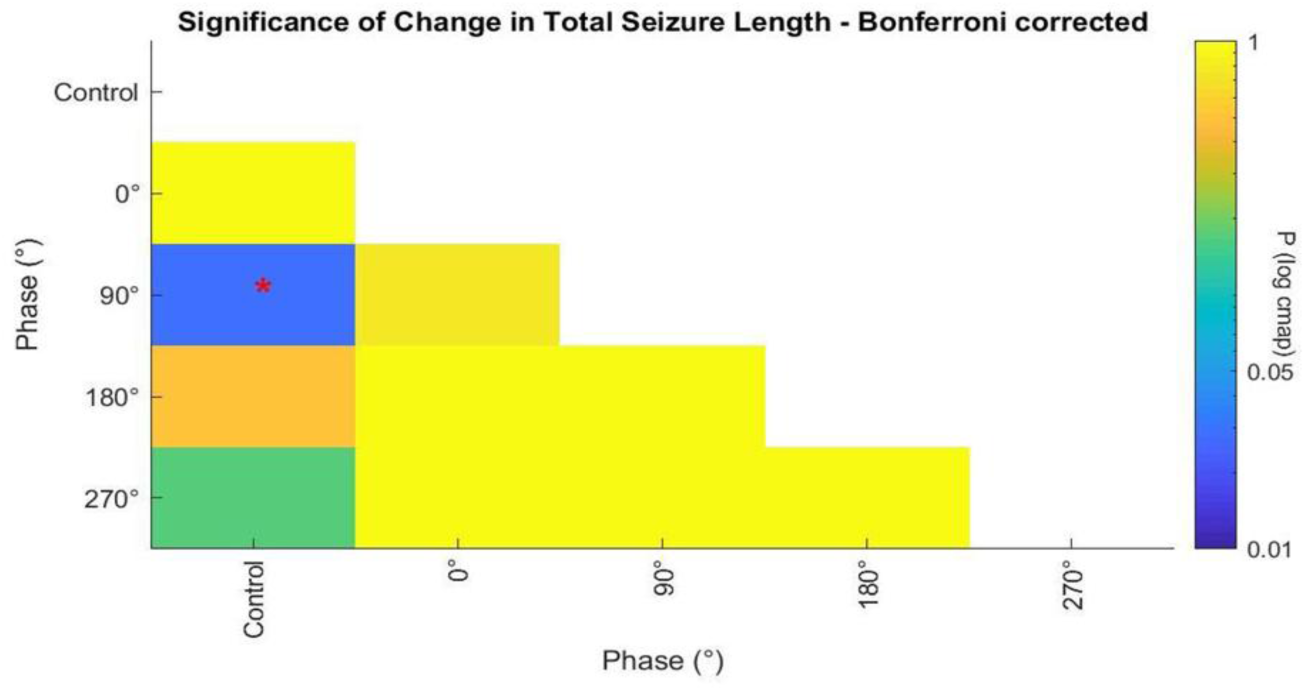
Statistical comparison of efficacy in various phase bins and control regarding the total seizure durations in humans, analyzed by the targeted ictal phase of the reconstructed deep-brain dipole at the seizure onset zone. The P-values of the Bonferroni-corrected pairwise comparisons of total seizure lengths are displayed on a logarithmic color scale for better visibility. Significant pairs (i.e. P < 0.05) are also marked with red asterisks for better readability.

**Figure S8.**
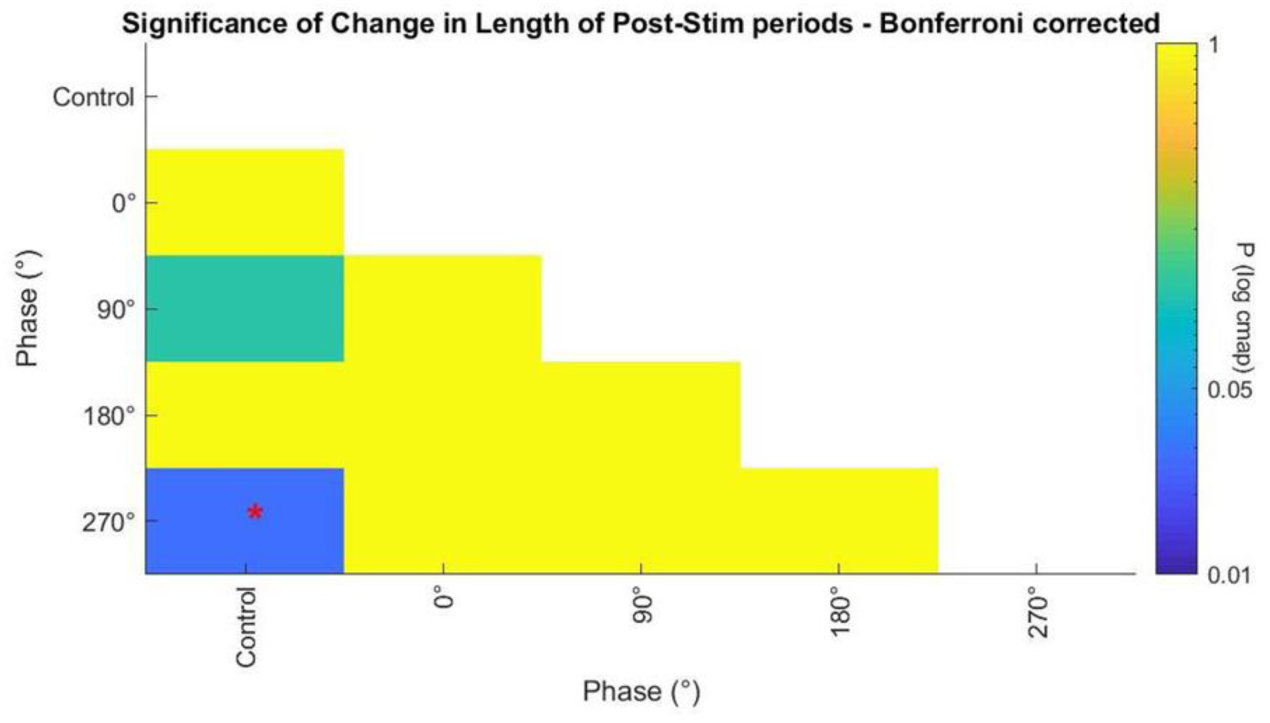
Statistical comparison of efficacy in various phase bins and control, regarding the post-stimulus seizure durations in humans, analyzed by the targeted ictal phase of the reconstructed deep-brain dipole at the seizure onset zone. The P-values of the Bonferroni-corrected pairwise comparisons of post-stimulation seizure lengths are displayed on a logarithmic color scale for better visibility. Significant pairs (i.e. P < 0.05) are also marked with red asterisks for better readability.

**Figure S9.**
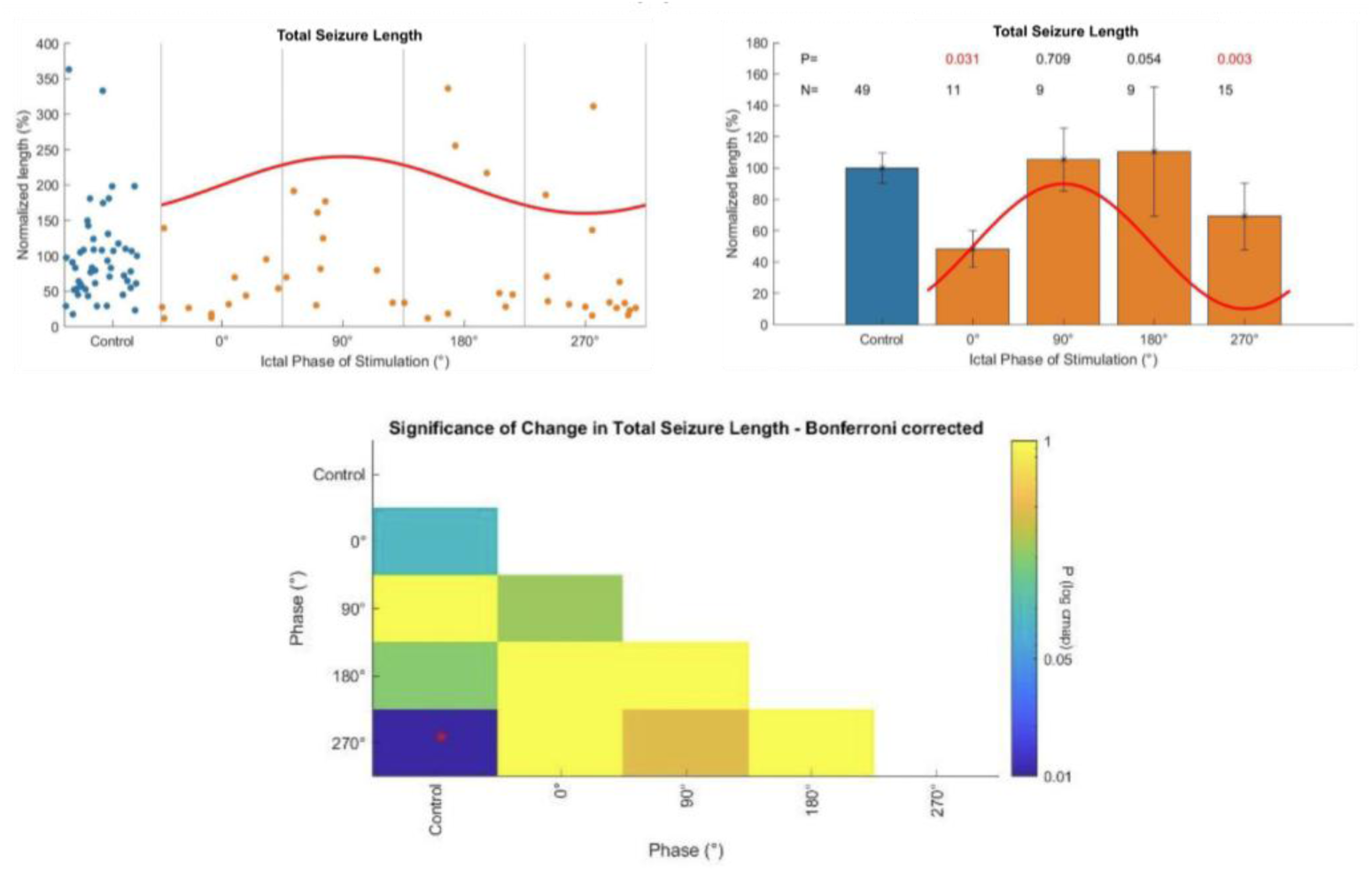
Efficacy of stimulation at various phase bins and control regarding the total seizure durations in humans, analyzed by the ictal phase of the surface EEG closest to the seizure onset zone. **(Top)** Change in total seizure duration as a function of the targeted ictal phase of the surface EEG. Data are normalized to each patient’s average control seizure length. Right panel shows the quantized measures (mean ± SEM) of the individual seizure data shown on the left panels as scatter plots. Unstimulated control cases are shown in blue. **(Bottom)** Pairwise comparison of the efficacy of various phase bins and control. The P-values of the Bonferroni-corrected pairwise comparisons are displayed on a logarithmic color scale for better visibility. Significant pairs (i.e. P < 0.05) are also marked with red asterisks for better readability.

**Figure S10.**
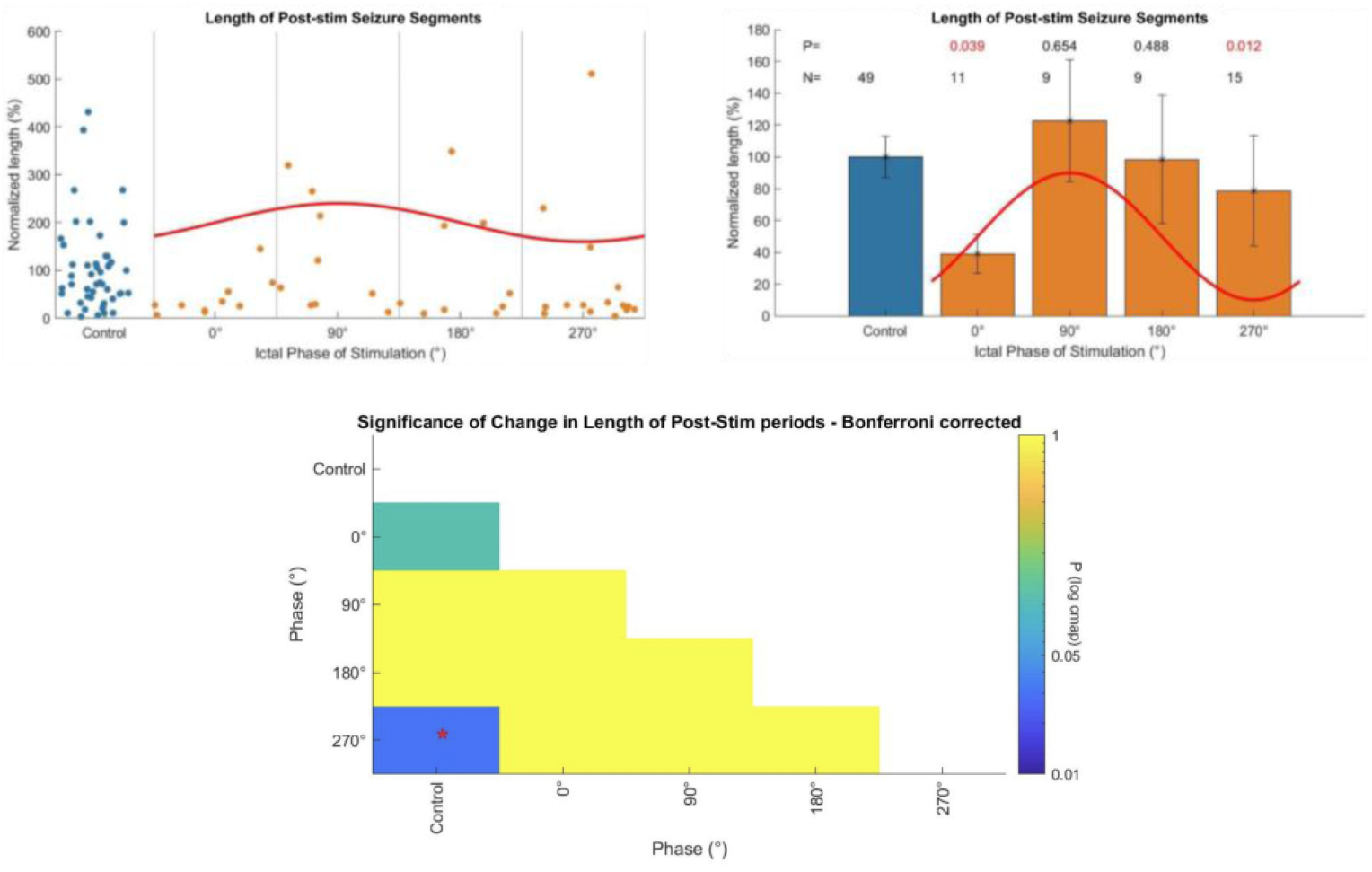
Efficacy of stimulation at various phase bins and control, regarding the post-stimulus seizure durations in humans, analyzed by the ictal phase of the surface EEG closest to the seizure onset zone. **(Top)** Change in remaining seizure duration after first stimulation, as a function of the targeted ictal phase of the surface EEG. Data are normalized to each patient’s average control seizure length. Right panel shows the quantized measures (mean ± SEM) of the individual seizure data shown on the left panels as scatter plots. Unstimulated control cases are shown in blue. **(Bottom)** Pairwise comparison of the efficacy of various phase bins and control. The P-values of the Bonferroni-corrected pairwise comparisons are displayed on a logarithmic color scale for better visibility. Significant pairs (i.e. P < 0.05) are also marked with red asterisks for better readability.

**Table S1.**
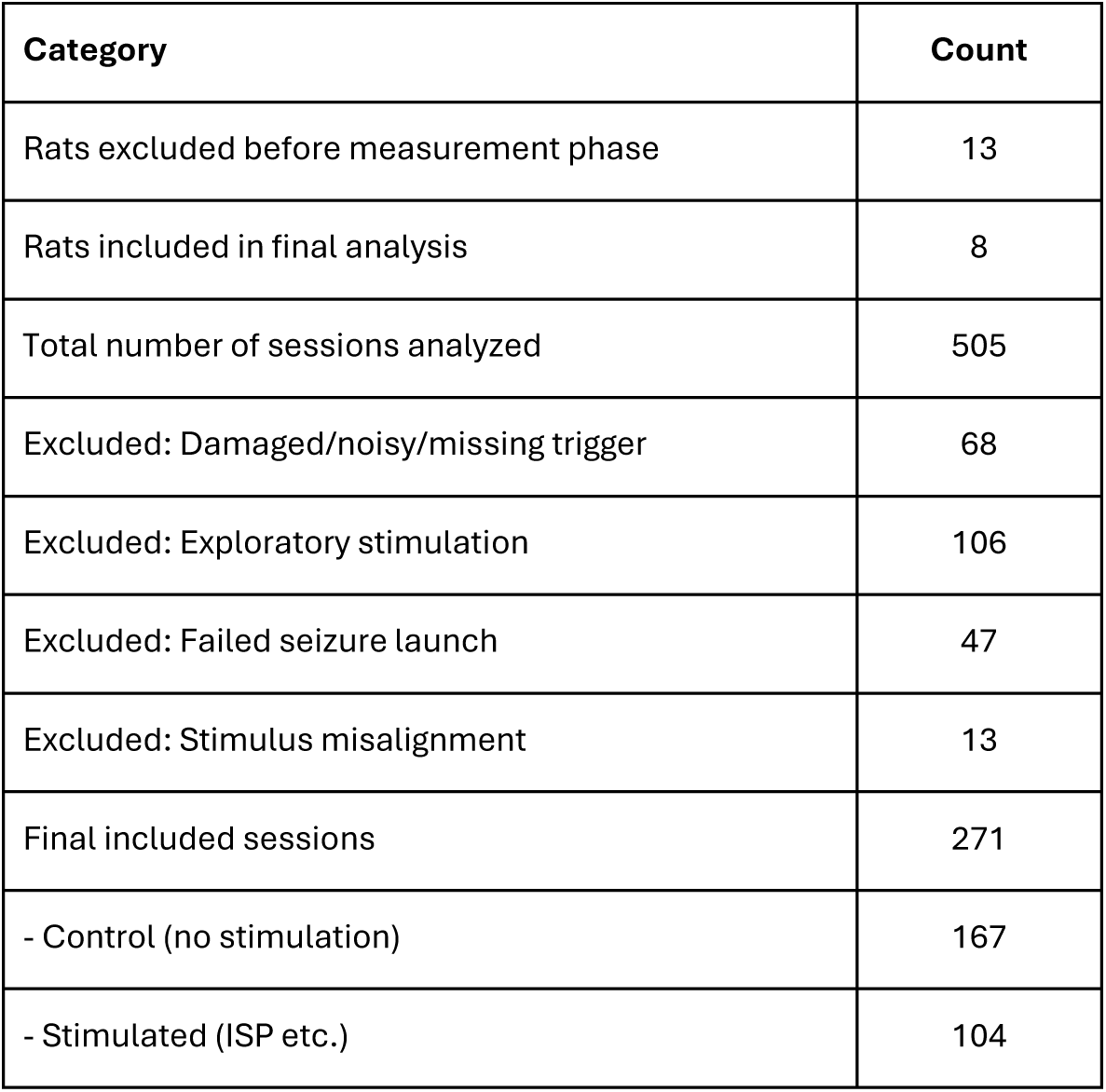
Overview of Animal Exclusion and Session Filtering During the Measurement Phase.

**Table S2.** Descriptive statistics and the results of the per animal comparison of total seizure length in rats (Kolmogorov- Smirnov tests)

| Animal ID |  | NSS07 | NSS11 | NSS12 | NSS14 | NSS15 | NSS17 | NSS20 | NSS21 |
| --- | --- | --- | --- | --- | --- | --- | --- | --- | --- |
| Control | Median (IQR) (s) | 107.72<br>(13.94) | 57.44<br>(62.48) | 65.64<br>(42.25) | 127.46<br>(35.4) | 113.53<br>(21.95) | 97.22<br>(33.16) | 97.02<br>(33.39) | 75.6<br>(29.2) |
|  | N | 13 | 51 | 53 | 9 | 16 | 11 | 8 | 6 |
| Stimulated | Median (IQR) (s) | 16.58<br>(9.62) | 48.03<br>(15.45) | 45.8<br>(26.47) | 69.97<br>(38.76) | 43.45<br>(71.8) | 68.08<br>(16.3) | 54.86<br>(46.4) | 55.23<br>(13.2) |
|  | N | 6 | 24 | 29 | 7 | 12 | 8 | 10 | 8 |
| P |  | 0.0024 | 0.0014 | 0.0005 | 0.0465 | 0.0021 | 0.0399 | 0.0473 | 0.1776 |

**Table S3.** Descriptive statistics and the results of the per animal comparison of the generalized segments in rats (Kolmogorov-Smirnov tests)

| Animal ID |  | NSS07 | NSS11 | NSS12 | NSS14 | NSS15 | NSS17 | NSS20 | NSS21 |
| --- | --- | --- | --- | --- | --- | --- | --- | --- | --- |
| Control | Median (IQR) (s) | 18.77<br>(12.86) | 29.51<br>(15.11) | 36.75<br>(18.71) | 28.04<br>(7.92) | 25.01<br>(15.96) | 23.74<br>(6.68) | 30.39<br>(7.37) | 38.36<br>(8.36) |
|  | N | 13 | 51 | 53 | 9 | 16 | 11 | 8 | 6 |
| Stimulated | Median (IQR) (s) | 6.3<br>(10.34) | 23.19<br>(8.95) | 24.91<br>(23.5) | 17.01<br>(12.73) | 21.75<br>(13.9) | 23.59<br>(16.47) | 20.29<br>(18.79) | 23.59<br>(19.45) |
|  | N | 6 | 24 | 29 | 7 | 12 | 8 | 10 | 8 |
| P |  | 0.0078 | 0.0127 | 0.0004 | 0.2763 | 0.437 | 0.601 | 0.0473 | 0.0797 |

**Table S4.** Descriptive statistics for full seizure durations and generalized seizure segments under ISP stimulation delivered at specific phases of the ongoing ictal oscillation in rats. Median seizure durations (% of control) are shown alongside interquartile ranges (IQR), sample sizes (N), and both uncorrected and Bonferroni-corrected p-values. Stimulation at 45° yielded the most pronounced seizure-suppressing effect in both seizure categories, with median durations reduced to 57.6% (full seizures) and 41.7% (generalized segments) of control values. Significant differences compared to control are marked in red for clarity. These results support the hypothesis that early-phase stimulation is the most effective in disrupting seizure dynamics.

| Metric |  | Control | 0° | 45° | 90°<br>(Peak) | 135° | 180° | 225° | 270°<br>(Trough) | 315° |
| --- | --- | --- | --- | --- | --- | --- | --- | --- | --- | --- |
| Full Seizure | Median (%) | 100.0 | 76.17 | 57.57 | 60.57 | 70.15 | 82.31 | 79.34 | 73.25 | 56.28 |
|  | IQR (%) | 55.2 | 31.02 | 26.08 | 70.88 | 34.67 | 62.01 | 20.19 | 33.98 | 26.11 |
|  | N | 167.0 | 15.0 | 12.0 | 7.0 | 20.0 | 12.0 | 6.0 | 10.0 | 22.0 |
|  | P |  | 0.001 | 0.0 | 0.0116 | 0.0001 | 0.162 | 0.059 | 0.002 | 0.0 |
|  | P (Bonf.) |  | 0.0049 | 0.0001 | 0.0348 | 0.0003 | 0.162 | 0.118 | 0.0078 | 3.1e-05 |
| Generalized Segments | Median (%) | 100.0 | 87.51 | 41.69 | 46.25 | 78.22 | 79.53 | 86.32 | 79.66 | 72.38 |
|  | IQR (%) | 36.35 | 30.85 | 36.09 | 30.66 | 38.35 | 34.31 | 22.12 | 32.53 | 40.68 |
|  | N | 167.0 | 15.0 | 12.0 | 7.0 | 20.0 | 12.0 | 6.0 | 10.0 | 22.0 |
|  | P |  | 0.199 | 0.0 | 0.0013 | 0.0041 | 0.1243 | 0.6321 | 0.0061 | 0.0032 |
|  | P (Bonf.) |  | 0.398 | 0.0004 | 0.0093 | 0.0203 | 0.373 | 0.6321 | 0.0246 | 0.0189 |

**Table S5.** Descriptive statistics and comparison of change in total seizure duration as a function of the targeted ictal phase of the reconstructed deep-brain dipole in humans.

| Phase | Mean Seizure Duration (%) | SEM (%) | N | P (vs. Control) | P (Bonferroni-corrected) |
| --- | --- | --- | --- | --- | --- |
| Control | 100.00 | 9.82 | 49 | — | — |
| 0° | 97.25 | 23.55 | 12 | 0.1794 | 0.1794 |
| 90° (Peak) | 39.14 | 5.89 | 8 | 0.0028 | **0.0113** |
| 180° | 91.03 | 25.64 | 14 | 0.0668 | 0.1336 |
| 270° (Trough) | 75.90 | 28.93 | 10 | 0.0204 | 0.0613 |

**Table S6.** Descriptive statistics and comparison of change in post-stimulation seizure duration, as a function of the targeted ictal phase of the reconstructed deep-brain dipole at the seizure onset zone in humans.

| Phase | Mean Seizure Duration (%) | SEM (%) | N | P (vs. Control) | P (Bonferroni-corrected) |
| --- | --- | --- | --- | --- | --- |
| Control | 100.00 | 13.17 | 49 | — | — |
| 0° | 114.26 | 34.96 | 12 | 0.3952 | 0.6516 |
| 90° (Peak) | 31.57 | 7.80 | 8 | 0.0138 | **0.0415** |
| 180° | 89.15 | 25.14 | 14 | 0.3258 | 0.6516 |
| 270° (Trough) | 73.10 | 48.79 | 10 | 0.0028 | **0.0113** |

**Table S7.** Descriptive statistics and comparison of change in total seizure duration as a function of the targeted ictal phase of surface EEG closest to the seizure onset zone in humans.

| Phase | Mean Seizure Duration (%) | SEM (%) | N | P (vs. Control) | P (Bonferroni-corrected) |
| --- | --- | --- | --- | --- | --- |
| Control | 100.00 | 9.82 | 49 |  |  |
| 0° | 48.25 | 11.91 | 11 | 0.0105 | **0.0314** |
| 90° (Peak) | 105.54 | 20.15 | 9 | 0.7092 | 0.7092 |
| 180° | 110.42 | 41.22 | 9 | 0.0268 | 0.0535 |
| 270° (Trough) | 69.28 | 21.24 | 15 | 0.0007 | **0.0027** |

**Table S8.** Descriptive statistics and comparison of change in post-stimulation seizure duration, as a function of the targeted ictal phase of surface EEG closest to the seizure onset zone in humans.

| Phase | Mean Seizure Duration (%) | SEM (%) | N | P (vs. Control) | P (Bonferroni-corrected) |
| --- | --- | --- | --- | --- | --- |
| Control | 100.00 | 9.82 | 49 |  |  |
| 0° | 39.22 | 12.22 | 11 | 0.0131 | **0.0393** |
| 90° (Peak) | 122.71 | 38.34 | 9 | 0.6536 | 0.6536 |
| 180° | 98.46 | 40.13 | 9 | 0.2438 | 0.4876 |
| 270° (Trough) | 78.72 | 34.68 | 15 | 0.0030 | **0.0121** |

